# Therapeutic role of recurrent ESR1-CCDC170 gene fusions in breast cancer endocrine resistance

**DOI:** 10.1101/2020.06.26.149948

**Authors:** Li Li, Ling Lin, Jamunarani Veeraraghavan, Yiheng Hu, Xian Wang, Sanghoon Lee, Ying Tan, Rachel Schiff, Xiao-Song Wang

## Abstract

**Background:** Endocrine therapy is the most common treatment for estrogen receptor (ER)-positive breast cancer, but its effectiveness is limited by high rates of primary and acquired resistance. There are likely many genetic causes and recent studies suggest the important role of *ESR1* mutations and fusions in endocrine resistance. Previously we reported a recurrent *ESR1* fusion called *ESR1-CCDC170* in 6-8% of the luminal B breast cancers that has a worse clinical outcome after endocrine therapy. Despite being the most frequent *ESR1* fusion, its functional role in endocrine resistance have not been studied *in vivo*, and the engaged mechanism and therapeutic relevance remain uncharacterized.

**Methods:** The endocrine sensitivities of HCC1428 or T47D breast cancer cells following genetic perturbations of ESR1-CCDC170 were assessed using clonogenic assays and/or xenograft mouse models. The underlying mechanisms were investigated by reverse phase protein array, western blotting, immunoprecipitation, and bimolecular fluorescence complementation assays. The sensitivity of ESR1-CCDC170 expressing breast cancer cells to concomitant treatments of tamoxifen and HER/SRC inhibitors was assessed by clonogenic assays.

**Results:** Our results suggested that different *ESR1-CCDC170* fusions endow different levels of reduced endocrine sensitivity *in vivo*, resulting in significant survival disadvantages. Further investigation revealed a novel mechanism that ESR1-CCDC170 binds to HER2/HER3/SRC and activates SRC/PI3K/AKT signaling. Silencing of ESR1-CCDC170 in the fusion-positive cell line, HCC1428, downregulates HER2/HER3, represses pSRC/pAKT, and improves endocrine sensitivity. More important, breast cancer cells expressing ectopic or endogenous ESR1-CCDC170 are highly sensitive to treatment regimens combining endocrine agents with the HER2 inhibitor lapatinib and/or the SRC inhibitor dasatinib.

**Conclusion:** ESR1-CCDC170 may endow breast cancer cell survival under endocrine therapy via maintaining/activating HER2/HER3/SRC/AKT signaling which implies a potential therapeutic strategy for managing these fusion positive tumors.

## Background

Endocrine therapy is the most effective treatment for estrogen-receptor (ER)-positive breast cancer (also known as luminal breast cancer). Agents targeting ER, including selective ER modulators (SERMs, i.e. Tamoxifen), selective ER down-regulators (SERDs, i.e. Fulvestrant) and aromatase inhibitors (AIs, i.e. letrozole) are the mainstays of treatment [1]. However, the efficacy of endocrine therapy is limited by intrinsic and acquired endocrine resistance [2]. About a quarter of the patients with primary tumor and almost all patients with metastases will present with or eventually develop endocrine resistance [3]. Tremendous efforts have been made to study the mechanism of endocrine resistance, and emerging evidence suggests that ESR1 mutations or fusions that mutate or eliminate its ligand binding domain constitute one of the most important driving mechanisms [3–6].

Recurrent gene fusions resulting from chromosome translocations are a critical class of genetic aberrations causing cancer [7], which have fueled modern cancer therapeutics. Recently, several milestone studies have identified recurrent gene fusions in different types of solid tumors with tremendous clinical impact. This is represented by the discovery of *EML4-ALK* fusion in ~4% of non-small cell lung cancer and *FGFR-TACC* fusion in ~3% of glioblastomas that have culminated in effective targeted therapies in these tumors [8, 9]. In particular, the discovery of EML4-ALK has led to accelerated approval of several ALK inhibitors by U.S. Food and Drug Administration (FDA) for the treatment of non-small cell lung cancer with stunning clinical responses [8]. Most recently, FDA granted accelerated approval to the first pan-cancer drug for the treatment of solid tumors, larotrectinib against the NTRK gene fusions [10]. Characterizing the role of gene fusions in breast cancer, particularly in endocrine resistance, will be critical for developing new and effective targeted therapies.

ER positive breast cancers can be classified into “luminal A” and “luminal B” subtypes. The luminal B breast tumors are more aggressive and endocrine-resistant luminal breast cancer that have high proliferative activity by Ki-67 index. Luminal B breast cancer accounts for 15-20% of all breast cancers [11], and is the most common subtype in young women [12]. In our previous study, through large-scale analyses of RNA-seq data from The Cancer Genome Atlas, we identified recurrent gene rearrangements between *ESR1* and its neighboring gene, coiled-coil domain containing 170 (*CCDC170*), in 6-8% of luminal B breast cancer, the majority of which are likely the result of tandem duplications [13]. Wild-type CCDC170 belongs to the structural maintenance of chromosome (SMC) protein family that maintains chromosome conformation through SMC-dependent looping and microtubule stabilization during interphase and mitosis [14, 15]. The *ESR1-CCDC170* fusions join the 5’ untranslated region of *ESR1* to the coding region of *CCDC170*, generating N-terminally truncated CCDC170 proteins (ΔCCDC170) expressed under the ESR1 promoter (Additional file 1: **Fig. S1**). This structure is distinct from other ESR1 fusions that retain its transcriptional activation domain [6]. When introduced into breast cancer cells, ESR1-CCDC170 proteins enhance ligand-independent growth factor signaling, leading to increased cell aggressiveness and tumorigenesis [13]. To date, ESR1-CCDC170 remains the most frequent gene fusion detected in luminal B breast cancer, and its recurrence has been subsequently supported by several recent studies [5, 16–19]. In addition, this fusion is also detected as a recurrent event in ovarian cancer, and its presence has been associated with exceptional short-term survival [20]. More interestingly, a recent publication reported the association of ESR1-CCDC170 fusions with lack of response to neoadjuvant letrozole treatment [17]. Nonetheless, detailed functional evidence demonstrating the role of ESR1-CCDC170 in endocrine resistance especially in the *in vivo* context, and the precise mechanism for this fusion to endow ligand-independent growth factor signaling has been lacking, which is not addressed in our previous study or the following reports. Furthermore, the therapeutic strategies to treat ESR1-CCDC170-positive tumors are ill-understood. Here we provide detailed evidence supporting the function of ESR1-CCDC170 in endowing breast cancer cell survival and reducing endocrine sensitivity *in vitro* and *in vivo*, and unravel its novel action mechanism via modulating and activating the SRC/HER2/HER3 complex. Our study suggests that the combination of inhibitors against these kinases with tamoxifen treatment as potential new therapeutic strategy for breast tumors harboring ESR1-CCDC170 fusions.

## Materials and Methods

### Cell culture

ER^+^ breast cancer cell lines T47D, HCC1428 and ZR-75-1 were purchased from American Type Culture Collection, cultured in RPMI-1640 (Corning) supplemented with 10% fetal bovine serum (Hyclone). For estrogen deprivation, cells were cultured in phenol red-free RPMI 1640 (Invitrogen) supplemented with 5% charcoal-dextran-stripped serum (Sigma).

### Engineering ectopic overexpression models

Lentiviral constructs and the lentivirus for ectopic expression of ESR1-CCDC170 fusion variants and wtCCDC170 were from our previous study [13]. The T47D cells were infected with lentivirus containing these constructs in medium containing 4μg/ml polybrene. Medium was replaced after overnight incubation. The cells with high GFP reporter expression were selected using flow cytometry two days later.

### Antibodies and Reagents

Primary antibodies against pEGFR-Y1068 (D7A5), EGFR, pHER2-Y877, pHER2-Y1221/1222 (6B12), pHER2-Y1248, HER2/ErbB2 (D8F12), pHER3-Y1289 (D1B5), HER3 (D22C5), pMet-Y1234/1235 (D26), Met (D1C2), pER-S118α (16J4), pERα-S167 (D1A3), ERα (D8H8), pAKT-S473 (D9E), AKT (C67E7), pERK (137F5), ERK (D13E1), pSrc-S416 (D49G4), Src (32G6), Integrin β1(D2E5) and Bcl-2 were purchased from Cell Signaling Technology. GFP antibody and HER2/neu Ab-8 (Clone e2-4001) mouse monoclonal antibody used for IP were obtained from Thermo Fisher. C6orf97 antibody is from GeneTex, V5 (MA5-15253) antibody is from Thermo Fisher and (Z)-4-Hydroxytamoxifen is from Sigma. Fulvestrant (ICI-182780, ZD 9238), Lapatinib (GW572016), Dasatinib, BEZ235 and AZD8931 were purchased from Selleckchem, Drugs were resolved in DMSO, diluted in culture medium before use.

### *In vivo* xenograft experiments

All animal experiments were performed in accordance with the institutional guidelines and regulations, and the animal protocol was approved by the BCM Institutional Animal Care and Use Committee (Approval # AN-6123). Briefly, 1×10^7^ T47D cells ectopically expressing the empty vector, or the ΔCCDC170 fusion variants E2-E7 and E2-E10 were resuspended in 20% Matrigel solution and injected bilaterally into 4-6 week-old female athymic nude mice (Harlan Sprague–Dawley), supplemented with 60-day-release 17β-estradiol pellets. Xenograft tumors of the T47D models were successfully engrafted in 6-9 mice per group. Growth of the xenograft tumors was monitored twice per week and tumor volume was measured using the formula 1/2 (length × width^2^). When the tumors reached 200 mm^3^, mice with tumors expressing the empty vector, E2-E7 or E2-E10 variants were randomized to +/− tamoxifen treatment. Tamoxifen (25 μg/kg body weight) was injected subcutaneously for 5 days/week. Growth of the xenograft tumors was monitored twice per week until the end of the experiment.

### siRNA and transfection

The E2-E10-specific siRNA (5’-CAUCACUGAGAUUAAAACU-3’), ERα-specific siRNA (5’-AGGCUCAUUCCAGCCACAGTT-3’), HER2-specific siRNA-1 (L-003126-00, ON-TARGETplus Human ERBB2 siRNA SMARTPool), HER2-specific siRNA-2 (5’-CACGUUUGAGUCCAUGCCCAA-3’), Src-specific siRNA-1 (J-003175-15, 5’-CCAAGGGCCUCAACGUGAA-3’) and Src-specific siRNA-2 (J-003175-16, 5’-GGGAGAACCUCUAGGCACA-3’) and control siRNA (D-001810-10-50, ON-TARGETplus Nontargeting Pool) were purchased from Dharmacon. All siRNAs were transfected using Lipofectamine RNAi MAX Reagent (Invitrogen) according to manufacturer’s instructions.

### Reverse phase protein array analysis

Reverse phase protein array assay was performed as described previously [21]. Briefly, protein lysates were prepared from HCC1428 cells with control or E2-E10 siRNA knockdown using modified Tissue Protein Extraction Reagent (TPER) (Pierce) and a cocktail of protease and phosphatase inhibitors (Roche Life Science). The protein lysates were then diluted into 0.5 mg/ml of total protein in SDS sample buffer and denatured on the same day. For each spot on the array, the background-subtracted foreground signal intensity was subtracted by the corresponding signal intensity of the negative control slide (omission of primary antibody) and then normalized to the corresponding signal intensity of total protein for that spot. The median of the triplicate experimental values (normalized signal intensity) is taken for each sample for subsequent statistical analysis.

### Protein subcellular localization

Fractionation of the nuclear and cytoplasmic proteins from T47D cells ectopically overexpressing the empty vector, wtCCDC170 or the ΔCCDC170 fusion variants E2-E7 or E2-E10 was performed using the NE-PER Nuclear and Cytoplasmic Extraction Kit (Thermo Fisher) according to manufacturer’s instructions. The fractionated protein samples (30□μgprotein) were then analyzed by western blot, as previously described [13].

### Immunoprecipitation

For IP with HER2/HER3/Src antibody. As described previously [22], cells were harvested and lysed in NETN-400 buffer (50 mM Tris-HCl, pH 8.0, 400 mM NaCl, 1 mM EDTA, and 0.5% Nonidet P-40) for 25 min on ice. The samples were centrifuged at 13,000 rpm for 15 min, and the supernatants were diluted with the same buffer without NaCl (NETN-0) to obtain a final concentration of NaCl at 150 mM. The samples were incubated with the appropriate antibodies at 4°C with rocking for 2 h. Protein G agarose was then added, and the incubation was continued for an additional 2 h. Beads were then washed three times using the NETN-150 buffer. The bound proteins were eluted with 100 mM glycine, pH 2.5, and then neutralized by adding 1/10 vol of 1 M Tris-Cl, pH 8.0. Eluted proteins were separated on 4–12% SDS-PAGE and blotted with the corresponding antibodies as indicated.

### Clonogenic assay

T47D cells were cultured in normal medium, RPMI1640 with phenol red and 10% FBS, and seeded at a density of 5000 cells per well in 24/48-well plate. After 24 hours, medium was replenished with normal medium or changed to ED medium, phenol red-free RPMI1640 supplemented with 5% charcoal-stripped serum (CSS), with or without indicated drugs in the presence or absence of siRNAs, for 14 days. For HCC1428, cells were seeded at a density of 5000 or 10000 cells per well in 24-well plate, with the indicated drugs added 24 hours later, and the cells were then cultured for another13 to 18 days. After that, the cells were fixed and stained with crystal violet, the colonies formed were calculated as intensity by using ImageJ software (National Institutes of Health) with ColonyArea plugin.

### Statistical analysis

The results of all *in vitro* experiments were compared by Student’s t-tests or Two-Way ANOVA, and all data are shown as mean ± standard deviation. For the *in vivo* study, statistical comparisons of tumor growth rates were performed using two-way mixed ANOVA that takes account of mice groups and time points as factors, and mouse subjects as random effects [23–25]. Long-term outcomes were evaluated by survival analysis methods. ‘Events’ were defined to mimic clinically relevant outcomes; time to tumor regression (tumorvolume-halving) was analyzed using Kaplan-Meier survival curves and compared by the generalized Wilcoxon test.

## Results

### *ESR1–CCDC170* fusions endow reduced endocrine sensitivity *in vitro* and *in vivo*

To explore the role of different forms of ESR1–CCDC170 fusions in endocrine resistance, we engineered four major fusion variants, E2-E6, E2-E7, E2-E8 and E2-E10, that join the exon 2 of *ESR1* with the exon 6, 7, 8 or 10 of *CCDC170*, respectively, into the T47D luminal breast cancer cells known to be dependent on oestrogen [26]. These fusion variants encode different sizes of N-terminally truncated CCDC170 (ΔCCDC170) proteins, which were verified by Western blotting (Additional file 2: **Fig. S2**). To explore its role in endocrine resistance, we assessed the consequences of ESR1-CCDC170 ectopic expression on the outcome of tamoxifen treatment *in vivo*. T47D xenograft tumors expressing vector, E2-E7, or E2-E10 fusion variants were developed in female athymic nude mice, implanted with estradiol (E2) pellets. When the tumors reached 150-200 mm^3^, the mice were randomized into tamoxifen treatment (25 ug/kg body weight) and untreated groups. In the absence of tamoxifen, tumor growth rates were significantly increased in the xenografts expressing E2-E10 fusion (P=0.001), and to a lesser degree in the xenografts expressing E2-E7 fusion (P=0.069, **Fig. 1A-B**). When treated with tamoxifen, the vector group showed true tumor regression with 51% decrease in size (P=0.000007 comparing +/−tamoxifen), whereas the E2-E7 expressing tumors sustained steady tumor volumes with only 14% decrease in size (P=0.003 comparing +/−tamoxifen). The E2-E10 expressing tumors continued to grow and then stabilized at higher tumor burdens, with almost two folds increase in tumor volumes (P=0.151 comparing +/− tamoxifen). The relative endocrine resistance of ESR1-CCDC170 expressing tumors was evident when the tumor growth rates of different models within the tamoxifen treated group are compared (vector vs E2-E7: P=0.001, vector vs E2-E10: P=0.000002, **Fig. 1B**). Kaplan–Meier analysis revealed a significantly worse regression-free survival for both E2-E7 (p<0.01) and E2-E10 (p<0.001)-overexpressing tumors treated with tamoxifen compared to the vector control tumors (**Fig. 1C**). These data suggest that *ESR1-CCDC170* variants render the T47D xenografts less sensitive to tamoxifen treatment in the *in vivo* context, and the E2-E7 and E2-E10 variants endow different levels of reduced responsivity.

**Figure 1.**
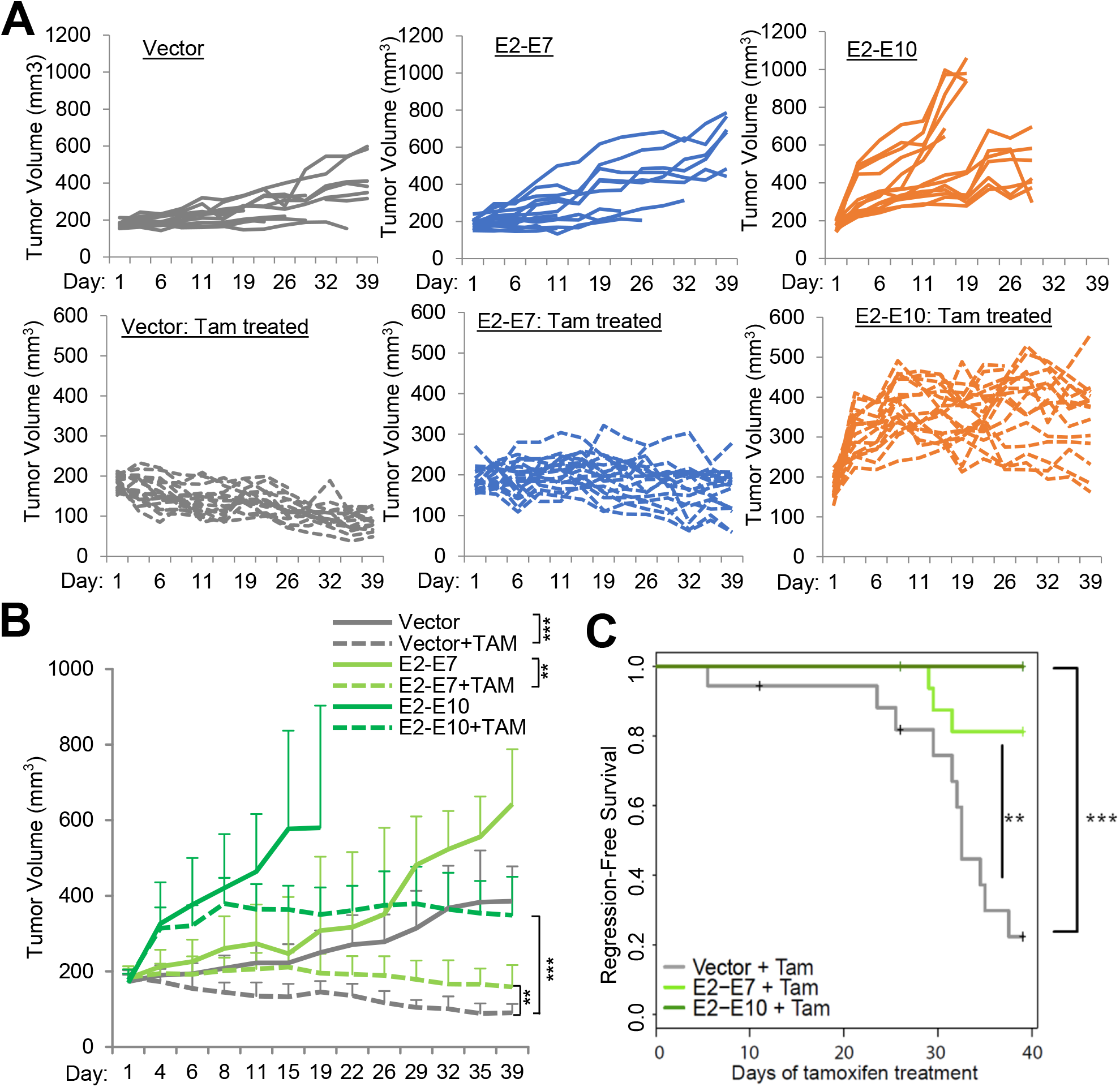
*ESR1–CCDC170* fusion variants endow tamoxifen resistance *in vivo*. **A)** The individual tumor growth curves of the T47D xenograft tumors expressing a vector, E2–E7 or E2–E10 *ESR1–CCDC170* fusion variants with or without tamoxifen treatment (TAM). Athymic female mice prepped with estrogen (E2) pellets were injected with 1×10^7^engineered T47D cells, and randomized into with/without tamoxifen treatment groups when the tumor volume reached 150-200 mm^3^. Vector, pLenti7.3 vector control. **B**) The summarized tumor growth curves of the T47D xenograft tumors treated with or without tamoxifen as in Fig. 1A. **P<0.01, ***P<0.001 (based on two-way mixed ANOVA). The statistical significance comparing +/− tamoxifen within each tumor model: Vector (P=0.000007), E2-E7 (P=0.003), E2-E10 (P=0.151). The statistical significance comparing different models within tamoxifen treated group are vector vs E2-E7 (P=0.001), vector vs E2-E10 (P=0.000002). **C**) the Kaplan–Meier curves for regression-free survival (tumor halving) of the engineered T47D xenograft tumors treated with tamoxifen. **P<0.01, ***P<0.001 (log-rank test).

Next, we assessed the effect of depleting ESR1-CCDC170 on the endocrine sensitivity of HCC1428, an ER^+^ breast cancer cell line harboring endogenous E2-E10 fusion. HCC1428 was derived from plural effusion of a 49 years old patient with metastatic luminal breast carcinoma after chemotherapy, who died 6 months later [27]. Thus, this cell line is endocrine-therapy naive. We used a validated siRNA targeting the E2-E10 fusion junction that can specifically knockdown the E2–E10 fusion, as we previously reported [13], and the silencing of E2-E10 protein was validated by western blot (Additional file 3: **Fig. S3A**). The cells were then treated with 4-hydroxytamoxifen (4-OHT), the active metabolite of tamoxifen used for *in vitro* experiments, or fulvestrant, a second-line endocrine agent that act by degrading ER. Tamoxifen or fulvestrant alone reduced the cell growth efficiently, with E2–E10 depletion led to more significant reduction of cell growth compared to the siRNA control (esp. in fulvestrant-treated cells, **Fig. 2A**), suggesting that silencing E2-E10 increases endocrine responsiveness of HCC1428 cells.

**Figure 2.**
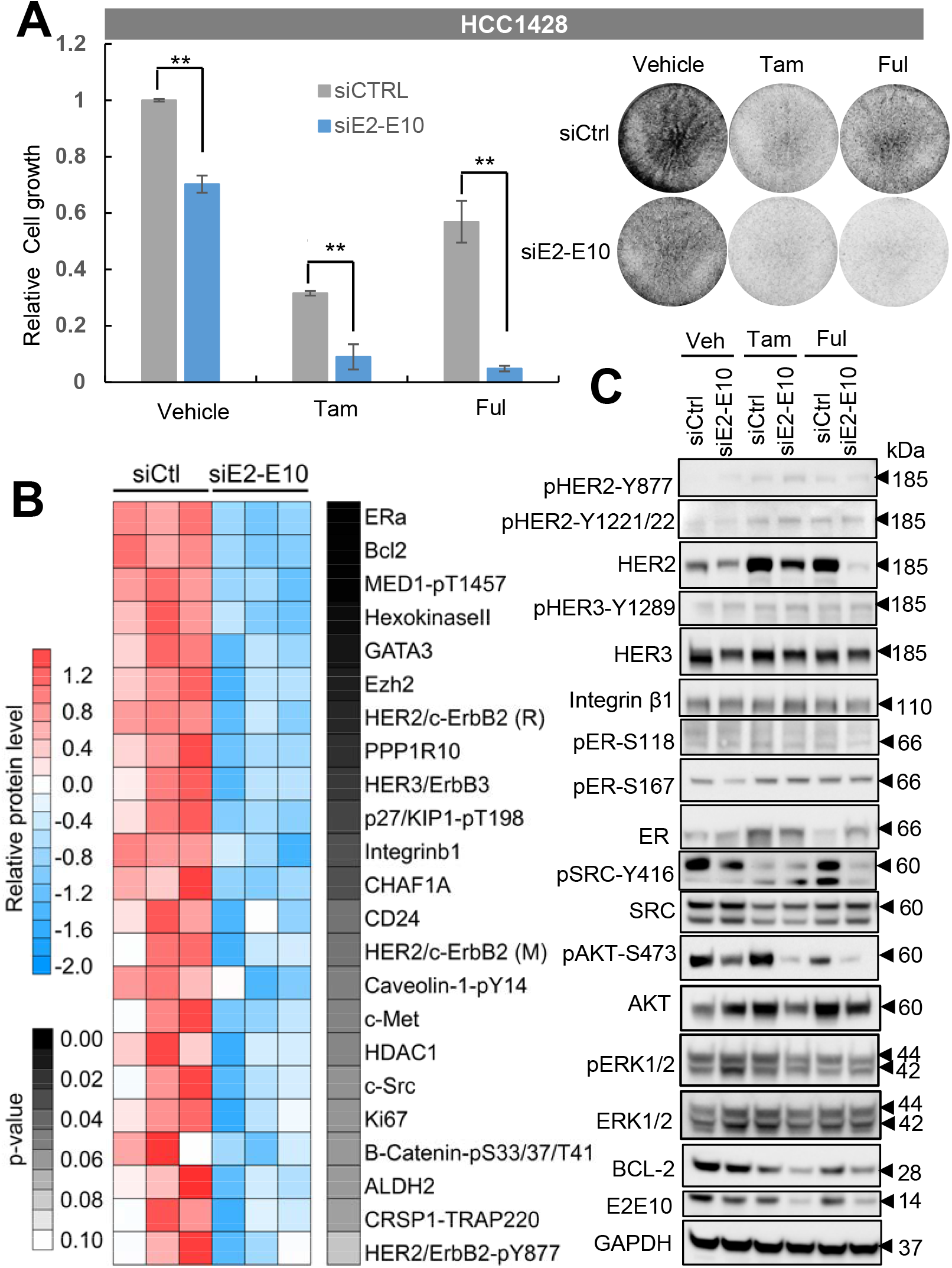
Silencing of *ESR1-CCDC170* increases the endocrine responsiveness of HCC1428 cells. **A)** Silencing of the E2-E10 fusion in HCC1428 cells reduced cell viability as shown by clonogenic assays. Cells were first treated with siRNAs for three days in biological triplicates, and then treated with 4-hydroxytamoxifen (Tam, 0.5 μM) or fulvestrant (Ful, 0.1 μM) for 72 hours, together with the siRNAs simultaneously. The left chart shows the relative intensity of triplicates (Means ± SD) normalized to the cells treated with vehicle and scramble siRNA control (siCtrl). The western blots verifying the knockdown efficiency and the representative images of clonogenic assays are shown on the right. **, *P*<0.01; Student’s t test. Experiments performed at least three times. **B)** Heatmap showing the top down regulated (p<0.1) signaling molecules in E2-E10-silenced HCC1428 cells revealed by RPPA data. HER2/c-ErbB2 (R) and HER2/c-ErbB2 (M) indicate rabbit and mouse antibodies detecting the total HER2 protein respectively. **C)** Protein extracts from the HCC1428 treated as in A) were analyzed by Western blot analysis of key signaling molecules revealed by RPPA.

### *ESR1–CCDC170* fusions augment HER2/HER3/SRC/AKT pathway

To gain insights into ESR1–CCDC170-driven mechanisms, we performed reverse phase protein array (RPPA) analysis in the HCC1428 cells with or without *ESR1-CCDC170* silencing, using about 200 validated antibodies against an array of cancer-related signaling molecules. Interestingly, ESR1-CCDC170 silencing leads to substantial repression of ERα, BCL2, HER3, c-SRC protein levels as well as total/phospho-HER2 (**Fig. 2B**). We further validated these alterations by Western blotting in E2-E10-silenced HCC1428 cells treated with vehicle, tamoxifen or fulvestrant. The protein levels of HER2, HER3, and BCL2 were indeed repressed following E2-E10 depletion. However, we did not observe repression of ERα protein level. Interestingly, HER2 protein level, but not transcript level, was upregulated by both tamoxifen and fulvestrant treatment, while depletion of E2-E10 counteracted this effect, especially under fulvestrant treatment (**Fig. 2C,** Additional file 3: **Fig. S3B**). In addition, pSRC-Y416 and pAKT-S473 were repressed following E2-E10 silencing in HCC1428 cells. These data suggest that E2-E10 may sustain cell survival via maintaining HER2/HER3 protein levels and activating SRC and AKT signaling in HCC1428 cells.

We then went on to assess the signaling alterations in T47D cells ectopically expressing ESR1-CCDC170 variants or wtCCDC170 following endocrine treatment. Interestingly, estrogen deprivation induced activation of pSRC-Y416, which was more potent in T47D expressing ESR1-CCDC170 variants compared to wtCCDC170 or vector control. Unlike pSRC-Y416, the activation of pHER3-Y1289 and pAKT-S473 were induced by estrogen deprivation in only E2-E8 or E2-E10 variant, respectively (**Fig. 3A**). When cells were treated with tamoxifen, increased total HER2/HER3 levels were observed in all engineered T47D cells. This is consistent with the previous report that tamoxifen can stimulate HER2/HER3 expression [28]. In addition, stronger activations of pHER2-Y1221/1222, pHER3-Y1289, pSRC-416 and pAKT-S473 were observed in T47D cells expressing the fusion variants compared to wtCCDC170 or vector control, suggesting that ESR1-CCDC170 fusions enhance the activation of HER2/HER3/SRC/PI3K/AKT signaling under tamoxifen treatment. Similar significant signaling alternations including increased activation of HER2, HER3, SRC, AKT, and ERK and increased EGFR, SRC and ER protein levels were observed following tamoxifen treatment in E2-E10-expressing T47D xenograft tumors, and to a lesser degree in E2-E7-expressing tumors (**Fig. 3C,** Additional file 4: **Fig. S4**), consistent with their tumor growth curves shown in **Fig. 1A-B**.

**Figure 3.**
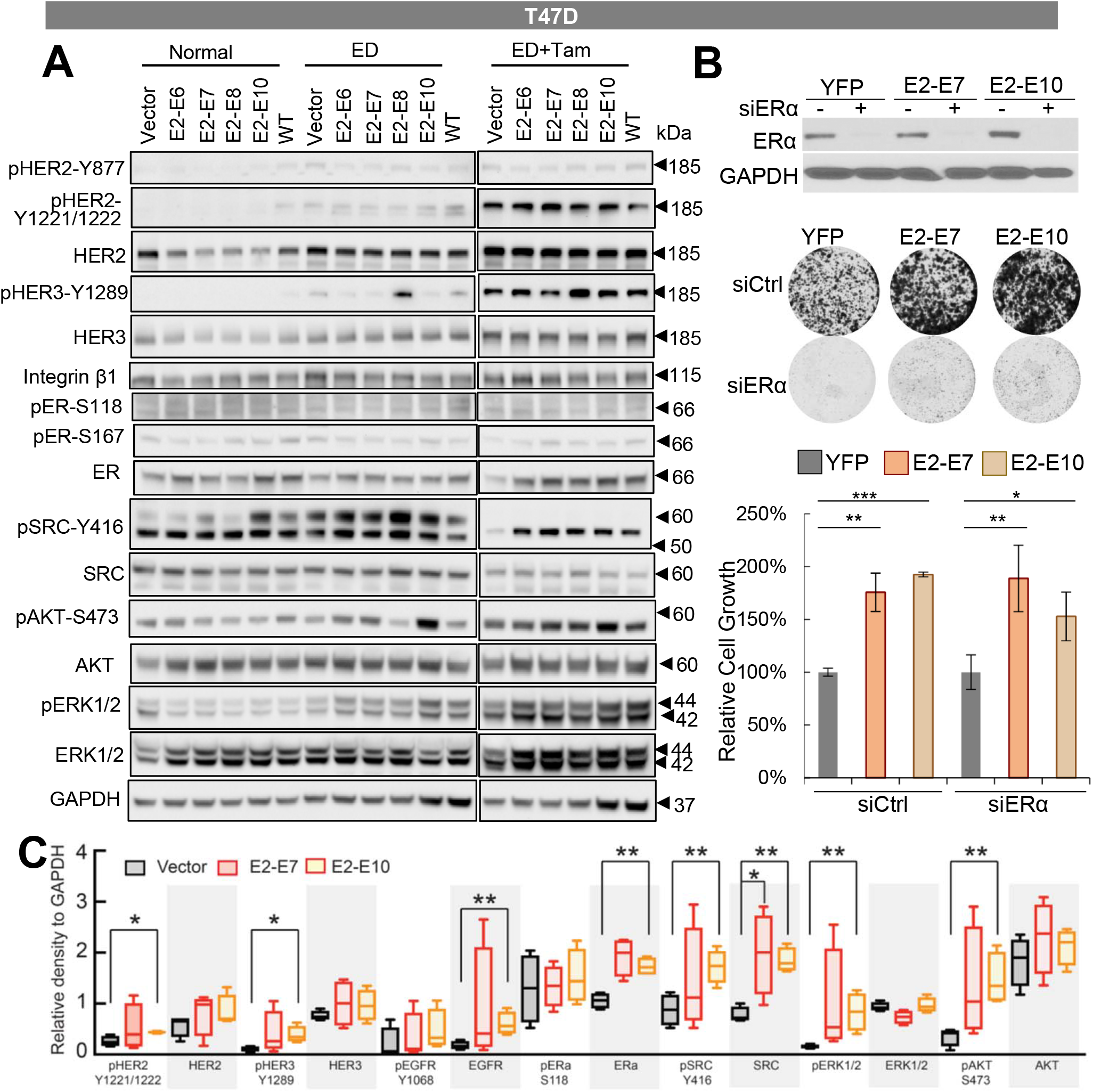
Signaling alterations in the engineered T47D cells following endocrine treatment *in vitro* and *in vivo*. **A**) Signaling alterations in the engineered T47D cells following endocrine treatment. Western blot analysis of the engineered T47D cells expressing *ESR1–CCDC170* fusion variants or wtCCDC170 cultured in normal medium (RPMI 1640 with phenol and 10% FBS), estrogen-deprived (ED) medium (phenol red-free RPMI 1640 with 5% CSS), or ED medium plus 4-OH tamoxifen (0.5 μM) for six days. **B**) The effect of ER depletion on the viability of engineered T47D cells expressing ESR1-CCDC170 variants or vector control as shown by clonogenic assays. Intensities of colonies in each well was normalized to the respective YFP group. The representative plate images are shown on the top. Upper panel, Western blots verifying the knockdown efficiencies of siERα. *P < 0.05, **P<0.01, ***P<0.001 (Student’s t-test). **C**) Western blot analysis of protein extracts from the engineered T47D xenograft tumor tissues treated with tamoxifen. Densitometric results of Western blots are shown in the figure. *p<0.05,**p<0.01 (Student’s t-test).

It is notable that the p-SRC-Y416 antibody detected two bands. The upper band that matches the predicted size of 60kD was diminished following tamoxifen treatment. SRC can be cleaved by calpain at N-terminal unique domain [29], which generates a truncated Src of 52kD, the size of the lower band. As calpain can be activated by endocrine treatment [30], it is possible that tamoxifen treatment may activate calpain, which then cleaves the 60kD SRC into a 52kD protein fragment. This suggests that the modulation of SRC under tamoxifen treatment is complicated, which requires further mechanism study to elucidate.

As many survival signaling pathways that drive endocrine resistance rely on modulating and reactivating ERα [31–33], we examined the effect of ESR1 depletion on the viability of engineered T47D cells expressing ESR1-CCDC170 variants or vector control. Clonogenic assays suggested that while ER depletion effectively inhibited cancer cell viability, ESR1-CCDC170 significantly increased the survival of T47D cells. This implies possible ER-independent mechanism engaged by ESR1-CCDC170 to endow cancer cell survival under endocrine therapy (**Fig. 3B**).

### *ESR1–CCDC170* localizes to cytoplasm, associates with HER2/HER3/SRC, and forms homodimers

Next, we asked whether ESR1-CCDC170 modulates the HER2/HER3/SRC pathway by forming complex with them. To test this, we performed immunoprecipitations using HER2, HER3, or SRC antibodies on the T47D cells ectopically expressing E2-E10 or wtCCDC170, and performed western blots using CCDC170 antibody. Interestingly, the ΔCCDC170 protein encoded by E2-E10 co-immunoprecipitated with endogenous HER2, HER3 and SRC (**Fig. 4A**), suggesting that E2-E10 forms complex with HER2, HER3 and SRC. On the other hand, only very modest wtCCDC170 protein bands were detected in the products immunoprecipitated with SRC/HER3, suggesting that the fusion protein has a much higher binding affinity to this complex. The interaction between HER2 and ΔCCDC170 was further verified on T47D cells ectopically expressing V5-E2-E7 or V5-E2-E10 via immunoprecipitation using V5 antibody and western blots using HER2 antibody (Additional file 5: **Fig. S5**). Next, we examined the subcellular localizations of ectopically expressed ESR1-CCDC170 proteins which revealed that the ΔCCDC170 proteins are more enriched in the cytoplasm, in contrast to the nuclear-enriched wtCCDC170 (Additional file 6: **Fig. S6**). Among the fusion variants, E2-E10 is mostly localized to cytoplasm, in line with its stronger effect on tamoxifen resistance.

**Figure 4.**
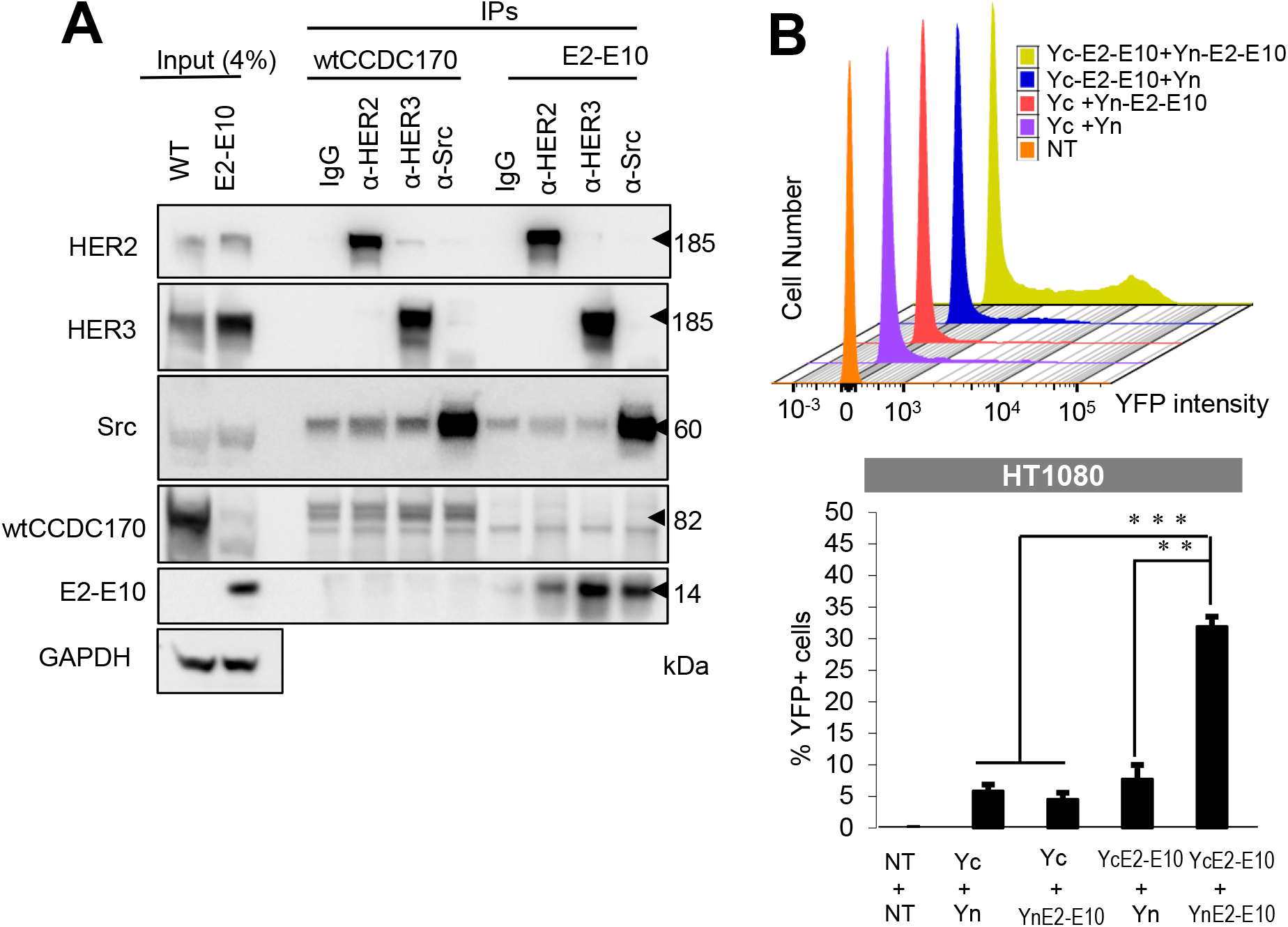
*ESR1–CCDC170* forms homodimers and interacts with HER2/HER3/SRC complex. **A)** ESR1-CCDC170 interacts with HER2, HER3 and SRC as shown by immuno-precipitation assay of T47D cells expressing wtCCDC170 or E2-E10, performed with anti-HER2, HER3, or SRC antibody, and detected by immuno-blotting with the indicated antibodies. **B)** BiFC assay to detect the dimerization of E2-E10 fusion protein. E2-E10 ORF was either fused to the N-terminal of YFP (Yn) or the C-terminal of YFP (Yc) for plasmid constructs. HT1080 cells were then co-transfected to express indicated Yn- and Yc-tagged proteins. The histogram shows the percentage of YFP-positive cells with the reconstituted YFP signal detected by flow cytometry in the respective transfected HT1080 cells.

We then investigated whether the N-terminal truncated CCDC170 proteins encoded by ESR1-CCDC170 fusions form dimers, which we speculate may help stabilize the HER2/HER3/SRC complex. To test this, we performed Bimolecular Fluorescence Complementation (BiFC) assay, which detects the proximity of two interacting proteins via reconstituted YFP fluorescence [34]. Here we tested the E2-E10 variant that encodes the smallest ΔCCDC170 protein with most N-terminal truncation. The E2-E10 open reading frame (ORF) was either fused to the N-terminal fragment of YFP (Yn) or to the C-terminal fragment of YFP (Yc), and then introduced into HT1080 cells, which possess the lowest auto-fluorescence, thus is most suited for BiFC assay [34, 35]. Western blots using GFP antibodies that cross-identify the YFP portion of proteins verified that Yn- or Yc-tagged E2-E10 was expressed at comparable levels between different transfected HT1080 cells (Additional file 7: **Fig. S7**). The reconstituted YFP signal was only detectable in the cells co-transfected with Yn-E2-E10 and Yc-E2-E10, suggesting that E2-E10 forms homodimer (**Fig. 4B**).

### SRC and HER2 inhibitors increase tamoxifen sensitivity in luminal breast cancer cells expressing *ESR1–CCDC170*

Next, we sought to test if targeting the HER or SRC signaling can revert the tamoxifen resistance driven by ESR1-CCDC170. We selected two drugs for this test, lapatinib, the first dual-specificity inhibitor targeting EGFR and HER-2 [36], and the FDA-approved SRC inhibitor, dasatinib [37, 38]. We cultured the T47D cells ectopically expressing ESR1-CCDC170 fusion variants in estrogen-deprived condition and treated them either with tamoxifen alone or combined with lapatinib or dasatinib. Clonogenic assay results showed that when treated with tamoxifen, T47D cells overexpressing the *ESR1–CCDC170* fusion variants grew more aggressively than that of the vector control and the wtCCDC170-expressing T47D cells, suggesting that *ESR1–CCDC170* fusion variants rendered T47D cells less dependent on oestrogen (**Fig. 5A**). Further, the colony formations were significantly reduced when the cells were treated with tamoxifen plus lapatinib (**Fig. 5A**) or dasatinib (**Fig. 5B**) compared to tamoxifen treatment alone, with dasatinib showing much more potent sensitizing effects.

**Figure 5.**
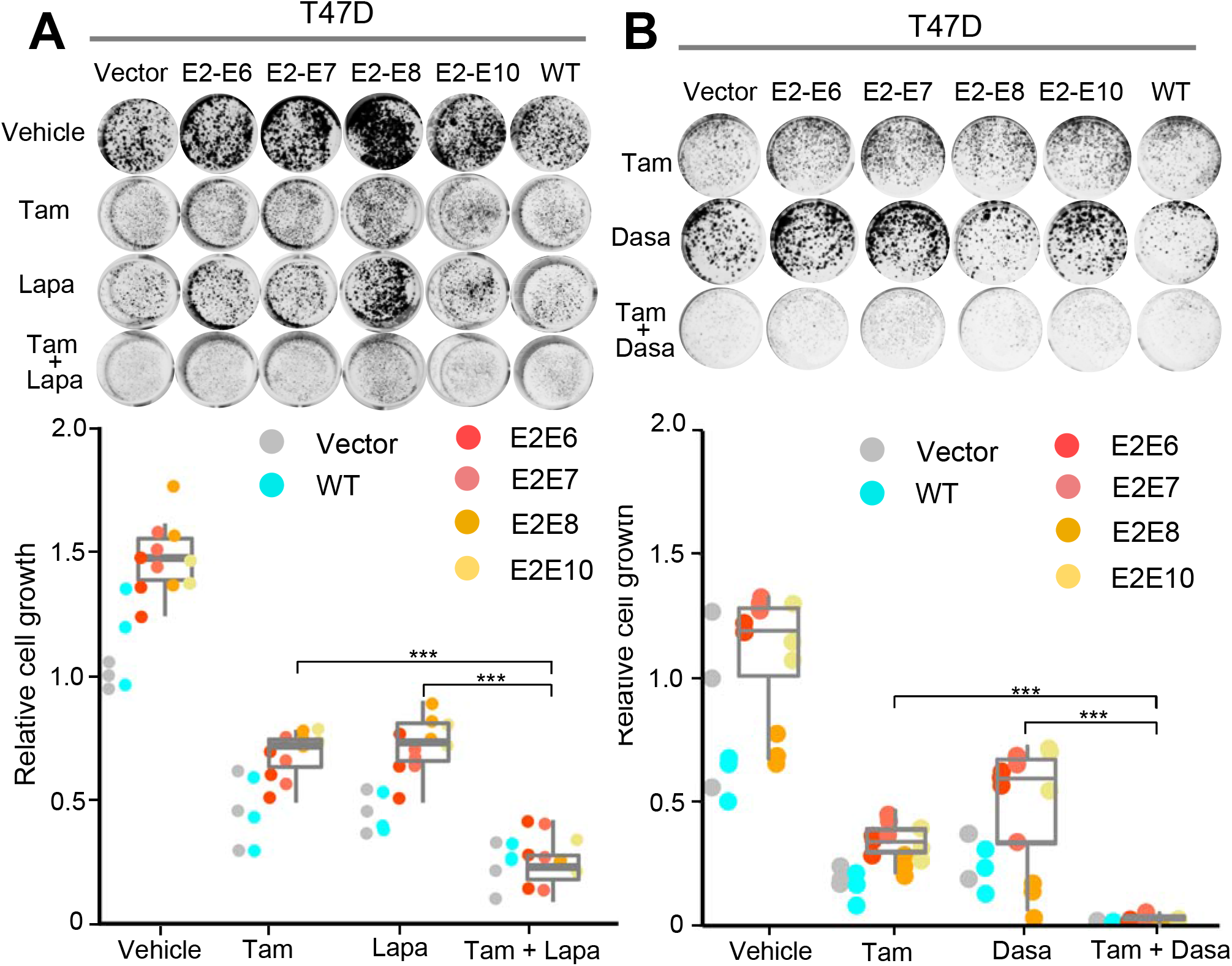
The response of engineered T47D cells expressing ectopic ESR1–CCDC17O fusion to HER2 or SRC inhibitors in combination with tamoxifen. **A)** The responses of the T47D cells expressing a control vector, ESR1–CCDC170 variants, or wtCCDC170 to tamoxifen (0.5 μM), lapatinib (0.5μM), or their combination as measured by clonogenic assays. The cells were cultured under phenol red-free medium and treated with the indicated drugs for 15 days before fixation and staining. Upper panel, the representative images of clonogenic assays. Lower panel, the relative intensities normalized to the vector control cells treated with vehicle (Means ± SD of triplicates). **B)** Responses of the engineered T47D cells to 4-OH tamoxifen (0.5 μM), SRC inhibitor dasatinib (0.05μM), or their combination as measured by clonogenic assay. The cells were cultured and treated in a similar way as in **A**. Upper panel, the representative images of clonogenic assays. Lower panel, the relative intensities normalized to the vector control treated with vehicle (Means ± SD of triplicates).

Next, we treated the HCC1428 cells expressing endogenous E2-E10 fusion with lapatinib, dasatinib, BEZ235 (PI3K/mTOR kinase inhibitor), or AZD8931 (a pan-ERBB inhibitor that inhibits EGFR, HER2, and HER3), in combination with tamoxifen or fulvestrant (**Fig. 6**). While all these inhibitors significantly inhibited cell viability compared to tamoxifen or fulvestrant treatment alone, lapatinib and dasatinib showed better therapeutic activities than BEZ235 or AZD8931. This suggests the importance of directly targeting SRC/HER2 to block fusion-driven signaling. In the presence of tamoxifen, dasatinib showed better activity than lapatinib (**Fig. 6A**), whereas when combined with fulvestrant, lapatinib and dasatinib showed comparable therapeutic effects. In addition, combining both lapatinib and dasatinib with fulvestrant resulted in additional therapeutic benefit which almost wiped out the cells (**Fig. 6B**). To verify the specific effect of HER2 or SRC inhibition in HCC1428 cells, we depleted HER2 or SRC using their specific siRNAs and treated the cells with tamoxifen or fulvestrant (**Fig. 7A-B**). The efficiency of these siRNAs was validated by western blots. Both HER2 and SRC silencing led to potent repression of cell viability especially when combined with fulvestrant treatment, similar to the effect of ESR1-CCDC170 silencing.

**Figure 6.**
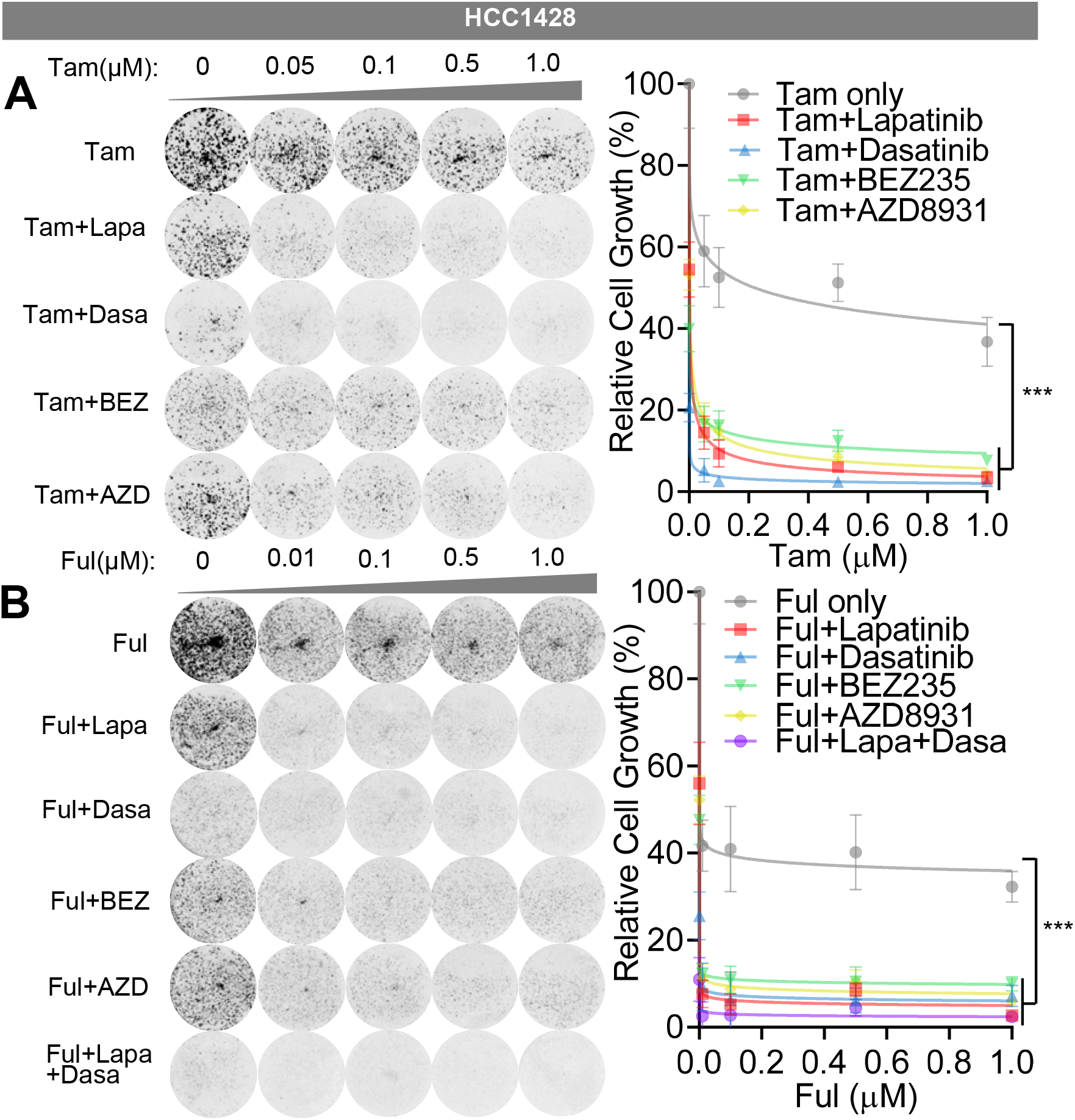
SRC and HER2 inhibitors increase endocrine sensitivity of HCC1428 cells expressing endogenous *ESR1–CCDC170* fusion. **A)** Sensitivity of HCC1428 cells to tamoxifen in combination with selective targeted agents as measured by clonogenic assays. HCC1428 cells were treated with different dosages of 4-OH tamoxifen (μM), or 4-OH tamoxifen plus lapatinib (2 μM), dasatinib (0.05 μM), BEZ235 (8 nM), or AZD8931 (1 uM). Data were normalized against vehicle treatment alone (as 100%). **B)** Sensitivity of HCC1428 cells to fulvestrant in combination with selective targeted agents as measured by clonogenic assays. HCC1428 cells were treated with different dosages of fulvestrant (μM), or fulvestrant plus lapatinib (2 μM), dasatinib (0.05 μM), BEZ235 (8 nM), AZD8931 (1 uM), or lapatinib (2 μM) + dasatinib (0.05 μM). Data were normalized against vehicle treatment alone (as 100%). Tam, 4-OH tamoxifen; Ful, fulvestrant, Lapa, lapatinib; Dasa, dasatinib, BEZ, BEZ235, AZD, ZAD8931. ***P < 0.001 (two-way ANOVA).

**Figure 7.**
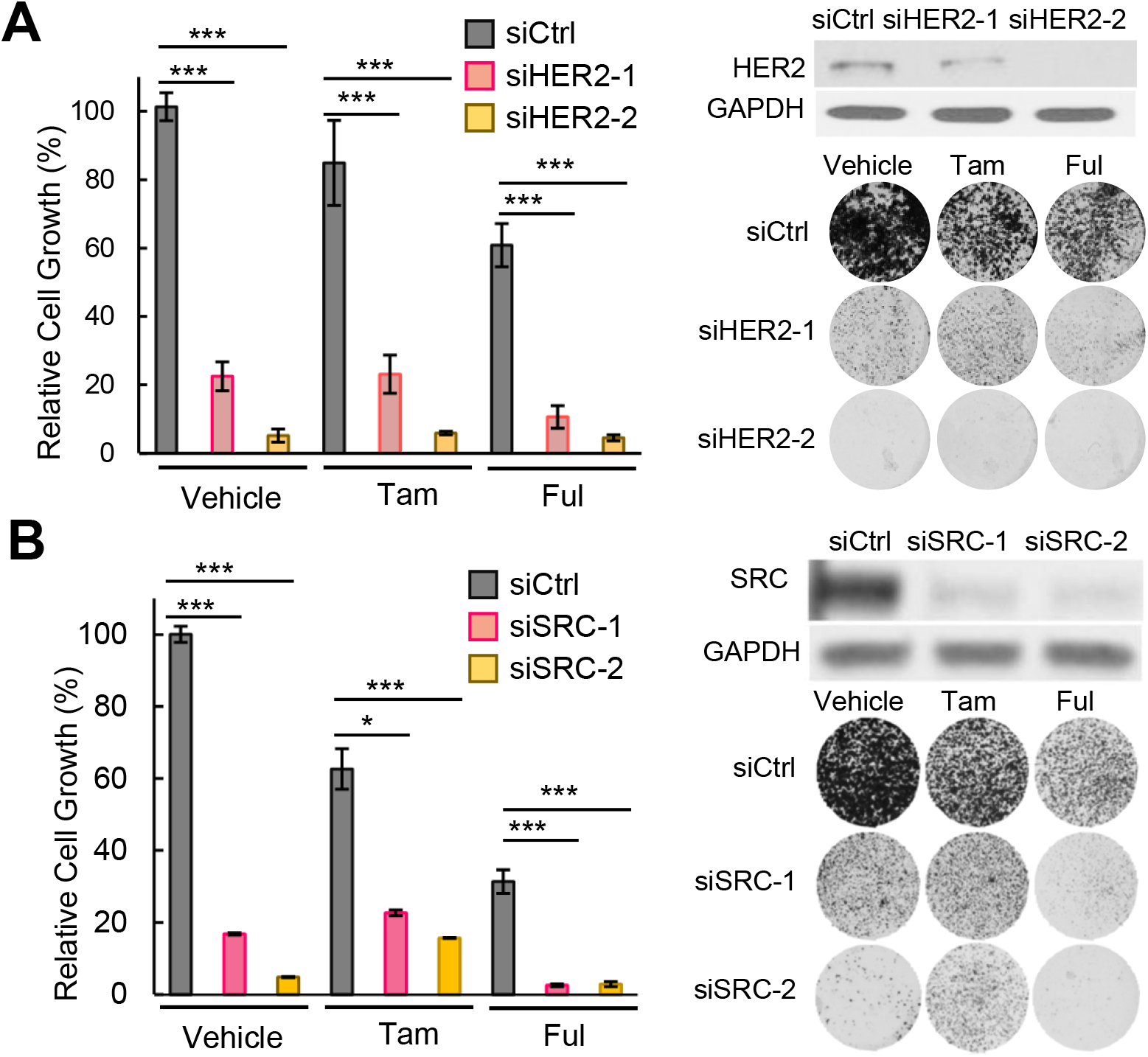
Silencing of HER2 or SRC increase endocrine responsiveness of HCC1428 cells. **A**) The effect of HER2 silencing on the endocrine responsiveness of HCC1428 cells as shown by clonogenic assays. Western blots verifying the knockdown efficiency and the representative plate images are shown on the right. Intensities of colonies in each well were normalized to the control siRNA and vehicle treated group. **B**) The effect of SRC silencing on the endocrine responsiveness of HCC1428 cells as shown by clonogenic assays. Western blots verifying the knockdown efficiency and the representative plate images are shown on the right. Intensities of colonies in each well were normalized to the control siRNA and vehicle treated group. Tam, 4-OH tamoxifen (0.5uM). Ful, Fulvestrant (0.1uM). ***P < 0.001, *P < 0.05 (student’s t-test).

Next, we assessed the therapeutic effect of lapatinib and dasatinib in ZR-75-1 cells harboring the E2-E6 fusion [13]. ZR-75-1 is an endocrine therapy-naïve cell line derived from ascitic effusion of a 63 year-old metastatic breast cancer patient who was subsequently treated with tamoxifen without apparent benefit [39]. This cell line was excluded from genetic silencing study as the fusion variant expressed in this cell line is not amendable to design siRNAs against its fusion junction due to their general toxicity to the cells. However, this cell line could be useful to provide additional insights into the therapeutic values of HER2/SRC inhibition in fusion positive cancer cells. We thus treated the ZR-75-1 cells with increasing doses of tamoxifen or fulvestrant, in combination with lapatinib, dasatinib, or both (Additional file 8: **Fig. S8A-B**). Our results showed that ZR-75-1 cells responded to both tamoxifen and fulvestrant treatment, with fulvestrant showing more potent effect. Concomitant lapatinib treatment resulted in further decreased cell viability compared to endocrine treatment alone, whereas dasatinib did not show any significant therapeutic benefit. Western blots revealed that the expression of SRC in ZR-75-1 cells appears to be at very low level (Additional file 9: **Fig. S9**). This suggests that the treatment strategy for managing ESR1-CCDC170 positive breast tumors may depend on the context of HER2 and SRC expressions which calls for further studies to elucidate.

## Discussion

Despite the tremendous success of endocrine therapies in ER^+^ breast cancer, endocrine resistance is a common and major clinical challenge. In this study, we provide molecular evidence supporting the role of ESR1-CCDC170 in breast cancer cell survival under endocrine therapy and the underlying mechanisms. Ectopic expression of the ESR1-CCDC170 fusion variants promote various levels of reduced endocrine sensitivities in the cell line and xenograft models. The relative tamoxifen-resistance of ESR1-CCDC170 expressing xenograft tumors are similar to that of the xenograft tumors expressing ectopic ESR1 mutations previously reported [40]. ESR1 mutations that mutate its ligand binding domain constitute one of the most important driving mechanisms of endocrine resistance [3, 4, 6], whereas the growth of the xenograft tumor models ectopically expressing ESR1 mutations can be effectively inhibited by endocrine treatment [40]. However, the relative endocrine resistance of the xenograft tumor models ectopically expressing ESR1 mutations was evident when the sizes of the different tumor models within the endocrine treated group are compared, similar to our observations [40]. Mechanistic studies suggest that ESR1-CCDC170 fusions form complex with HER2/HER3/SRC, augment HER2/HER3 protein levels and signaling, and enhance the activation of SRC/PI3K/AKT signaling under endocrine therapy *in vitro* and *in vivo*.

Overexpression or hyper-activation of the HER/SRC tyrosine receptor kinase family is known to contribute to endocrine resistance in breast cancer [2, 41–43]. HER2 and HER3 form heterodimer which is dependent on the HER3 ligand, and is known to exert potent mitogenic signal [44]. HER2/HER3 heterodimer functions as a major oncogenic unit and is known to crosstalk with SRC and activates PI3K/AKT pathway to drive breast cancer [36, 45]. SRC is broadly overexpressed in luminal breast cancer [46], and can crosstalk with HER2 when facilitated by other molecules such as CDCP1 [47]. As a result, SRC promotes the phosphorylation and activation of HER2, which in turn activates SRC [47]. Further, SRC activation has been reported to endow endocrine resistance through phosphorylating ER [31–33]. It is possible that the autonomous dimerization of ESR1-CCDC170 may help stabilize the interactions between HER2/HER3/SRC tyrosine kinases, and thus enhance their downstream signaling. Future studies will be required to pinpoint the precise mechanisms by which ESR1-CCDC170 fusions regulate HER2/HER3/SRC signaling.

Wild type CCDC170 contains a structural maintenance of chromosomes (SMC) domain (Additional file 1: **Fig. S1**). SMC proteins are formed by two long coiled-coil domains connected by a non-helical sequence called ‘hinge’ which presumably corresponds to the low complexity region of CCDC170. The SMC proteins contain highly conserved ATP-binding cassette (ABC) that drives its dimerization, and fold back on themselves at the hinge region through antiparallel coiled-coil interactions [48]. The ΔCCDC170 proteins encoded by the ESR1-CCDC170 fusions have different degrees of deletion of the N-terminal region of the SMC domain, but retain a putative high-affinity ATP-binding pocket at C-terminus. The N-terminal truncations of CCDC170 may expose the coiled-coil domain via reducing antiparallel coiled-coil interactions and thus alter its interactome, which calls for further investigations.

It should be stressed that while ESR1-CCDC170 functions through modulating SRC/HER2/HER3 complex, we cannot rule out the possibility that ESR1-CCDC170 could engage ERα to promote endocrine resistance. In fact, many survival signaling that drive endocrine resistance are known to act through phosphorylating and reactivating ER [31–33]. For example, SRC and AKT are known to phosphorylate Y537 and S167 respectively to modulate ER activity and endow endocrine resistance [31–33]. Thus, the survival signaling for endocrine resistance is not expected to act completely independent of ER. Future studies will be required to elucidate the function of ESR1-CCDC170 in modulating ERα activity under endocrine stress.

Furthermore, this study suggests a potential strategy to manage ESR1-CCDC170 positive patients via combining the HER2 inhibitor lapatinib and/or SRC inhibitor dasatinib with endocrine therapy. HER2 is amplified in approximately 10-15% of luminal breast cancer. While modest HER2 protein expression (IHC 1+ or 2+) can be detected in 60% of luminal breast cancer [49], HER2 inhibitors are generally not indicated in these tumors. While SRC has been reported to play a key role in breast cancer bone metastasis and hormonal therapy resistance [50–52], the clinical trials of SRC inhibitors in unselected metastatic breast cancer patients have been disappointing [50]. Our *in vitro* therapeutic studies on the T47D, HCC1428, and ZR-75-1 models suggest that the patients may be treated with HER2 and/or SRC inhibitors in combination with endocrine therapy depending on the context of HER2 and SRC expression levels. Since Her2 expression can be induced by endocrine treatment, but not SRC (**Fig. 2C, 3A**), the base-line expression of SRC could be particularly important for determining the therapeutic value of SRC inhibitor. Future preclinical and clinical studies are warranted to elucidate the optimal therapeutic strategy to manage ESR1-CCDC170-positive patients in the clinical setting.

## Conclusions

This study provides new molecular and functional evidence supporting the role of ESR1-CCDC170 in reducing endocrine responsiveness in breast cancer cells *in vitro* and *in vivo*, and revealed a novel action mechanism that ESR1-CCDC170 binds to HER2/HER3/SRC complex and activates their downstream signaling to endow cancer cell survival under endocrine therapy. Our results also imply a potential therapeutic strategy to manage ESR1-CCDC170-positive breast cancer patients via combining HER2 and/or SRC inhibitors with endocrine therapy.

## Additional Files

**Additional File 1: Figure S1.** Schematic of the ESR1-CCDC170 fusion variants and the encoded proteins.

**Additional File 2: Figure S2.** Verifying the ectopic expression of ESR1-CCDC170 fusion protein products in engineered T47D cells lines by Western Blots.

**Additional File 3: Figure S3.** Verifying the efficiency of E2-E10 siRNA in HCC1428 cells and its effect on ESR1 and HER2 mRNA expression.

**Additional File 4: Figure S4.** Western blot analysis of protein extracts from the engineered T47D xenograft tumor tissues treated with tamoxifen shown in Fig. 1A.

**Additional File 5: Figure S5.** Ectopically expressed V5-tagged ΔCCDC170 co-precipitates with HER2.

**Additional File 6: Figure S6.** Subcellular localization of ESR1-CCDC170 and wtCCDC170 proteins in the engineered T47D cells.

**Additional File 7: Figure S7.** Western blots related to Fig. 4B.

**Additional File 8: Figure S8.** The effect of SRC and HER2 inhibitors on the endocrine response of ZR-75-1 cells expressing endogenous ESR1–CCDC170 fusion.

**Additional File 9: Figure S9.** Western blots detecting HER2, HER3, and SRC protein expression in the cell models used in this study.

**Additional File 10:** The normalized RPPA data generated in this study.

## Abbreviations

ABC: ATP-binding cassette
AIs: aromatase inhibitors
BiFC: Bimolecular Fluorescence Complementation
CCDC170: coiled-coil domain containing 170
CSS: charcoal-stripped serum
ΔCCDC170: truncated CCDC170 proteins
E2: estradiol
ER: estrogen receptor
FDA: Food and Drug Administration
4-OHT: 4-hydroxytamoxifen
ORF: open reading frame
RPPA: Reverse phase protein array analysis
SERDs: selective ER down-regulators
SERMs: selective ER modulators
siRNA: small interfering RNA
SMC: structural maintenance of chromosomes
TPER: Tissue Protein Extraction Reagent

## Acknowledgements

We thank Dr. Shixia Huang, Hsin-Yi Cincy Lu, and Mr. Carlos Ramos from the Antibody-based Proteomics Core/Shared Resource for their excellent technical assistant in performing RPPA experiments. We thank Drs. Kimal Rajapakshe and Cristian Coarfa, Mr. Dimuthu Perera for RPPA data processing and normalization. We thank Suet Kee Loo for grammatical edits.

## Authors’ contributions

Li Li. designed and performed cell biology, mechanistic experiments, analyzed the data and wrote the manuscript. Ling Lin performed cell biology and mechanistic studies. J.V. performed the xenograft mouse and E2-E10 knockdown experiments. Y.H. performed fractionation assay and the BiFc experiments. X.W. performed the V5-IP experiments. S.L. analyzed the data. Y.T. generated the plasmid of the fusion variants. R.S. provided advices on the study and manuscript. X.-S.W. conceived and supervised the study, analyzed the data, and wrote the manuscript.

## Availability of data and materials

All data generated or analyzed during this study are included in this published article and its supplementary information files.

## Ethics approval and consent to participate

All animal experiments were performed in accordance with the institutional guidelines and regulations, and the animal protocol was approved by the BCM Institutional Animal Care and Use Committee (Approval # AN-6123).

## Consent for publication

Not applicable.

