## Additional File 1 for "Therapeutic role of recurrent ESR1-CCDC170 gene fusions in breast cancer endocrine resistance"

### Slide 1
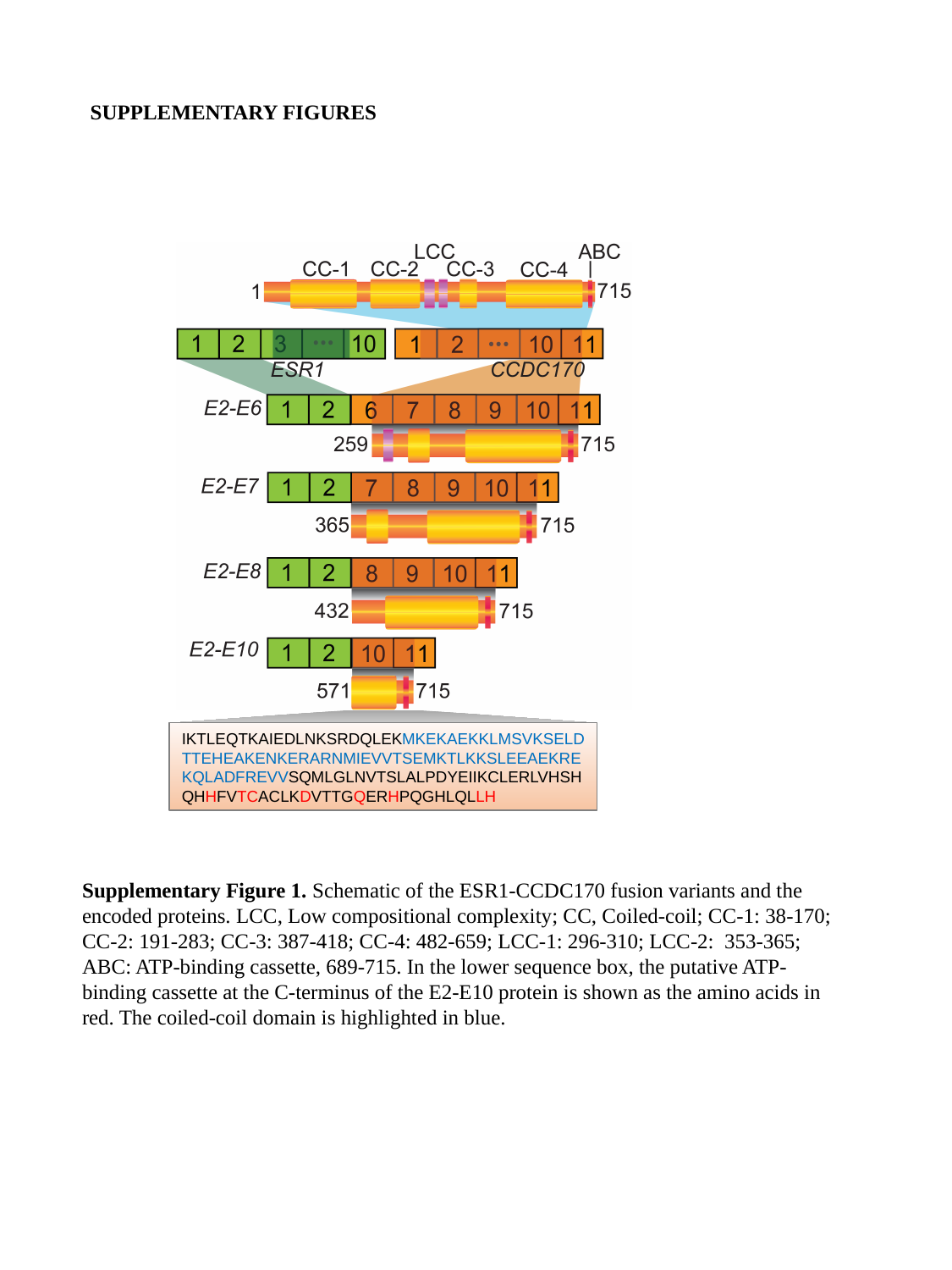

SUPPLEMENTARY FIGURES
IKTLEQTKAIEDLNKSRDQLEKMKEKAEKKLMSVKSELDTTEHEAKENKERARNMIEVVTSEMKTLKKSLEEAEKREKQLADFREVVSQMLGLNVTSLALPDYEIIKCLERLVHSHQHHFVTCACLKDVTTGQERHPQGHLQLLH
Supplementary Figure 1. Schematic of the ESR1-CCDC170 fusion variants and the encoded proteins. LCC, Low compositional complexity; CC, Coiled-coil; CC-1: 38-170; CC-2: 191-283; CC-3: 387-418; CC-4: 482-659; LCC-1: 296-310; LCC-2: 353-365; ABC: ATP-binding cassette, 689-715. In the lower sequence box, the putative ATP-binding cassette at the C-terminus of the E2-E10 protein is shown as the amino acids in red. The coiled-coil domain is highlighted in blue.
