## Additional File 2 for "Therapeutic role of recurrent ESR1-CCDC170 gene fusions in breast cancer endocrine resistance"

### Slide 1
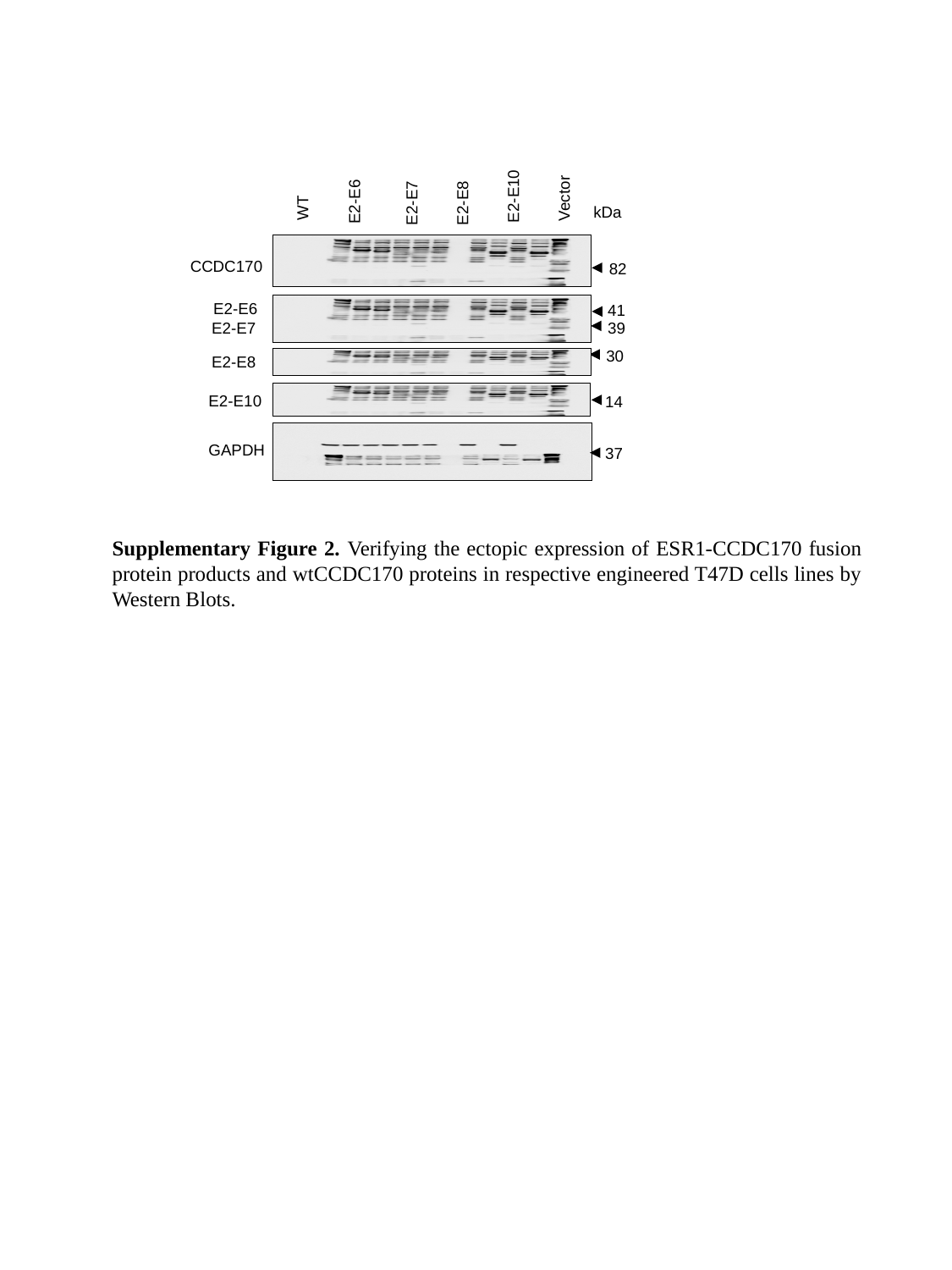

E2-E10
Vector
E2-E6
E2-E7
E2-E8
WT
CCDC170
E2-E6
E2-E7
E2-E8
E2-E10
GAPDH
kDa
82
41
39
30
14
37
Supplementary Figure 2. Verifying the ectopic expression of ESR1-CCDC170 fusion protein products and wtCCDC170 proteins in respective engineered T47D cells lines by Western Blots.
