## Additional File 3 for "Therapeutic role of recurrent ESR1-CCDC170 gene fusions in breast cancer endocrine resistance"

### Slide 1
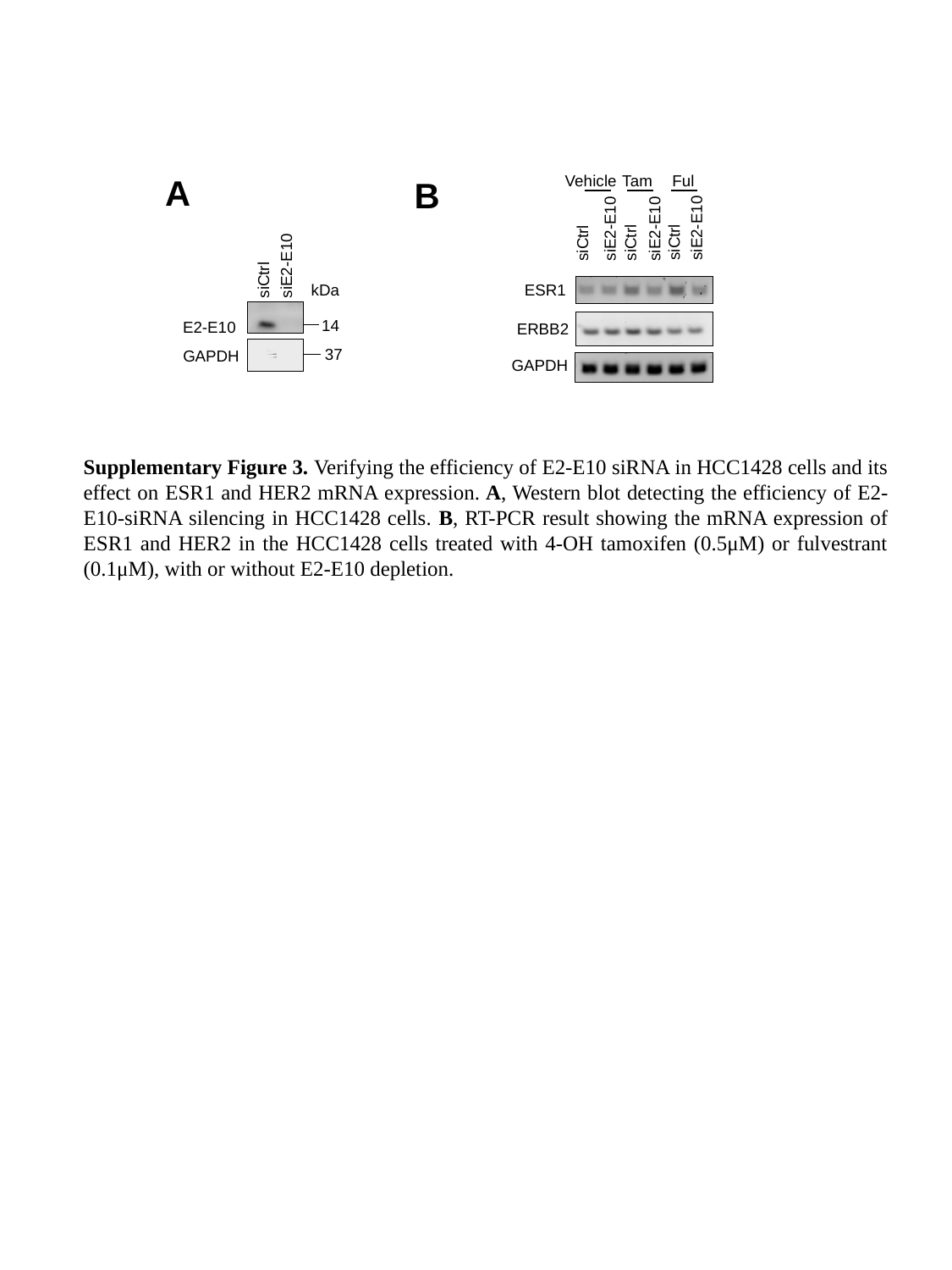

Tam
Ful
Vehicle
siE2-E10
siE2-E10
siE2-E10
siCtrl
siCtrl
siCtrl
ESR1
ERBB2
GAPDH
A
B
siE2-E10
siCtrl
kDa
14
E2-E10
37
GAPDH
Supplementary Figure 3. Verifying the efficiency of E2-E10 siRNA in HCC1428 cells and its effect on ESR1 and HER2 mRNA expression. A, Western blot detecting the efficiency of E2-E10-siRNA silencing in HCC1428 cells. B, RT-PCR result showing the mRNA expression of ESR1 and HER2 in the HCC1428 cells treated with 4-OH tamoxifen (0.5μM) or fulvestrant (0.1μM), with or without E2-E10 depletion.
