## Additional File 4 for "Therapeutic role of recurrent ESR1-CCDC170 gene fusions in breast cancer endocrine resistance"

### Slide 1
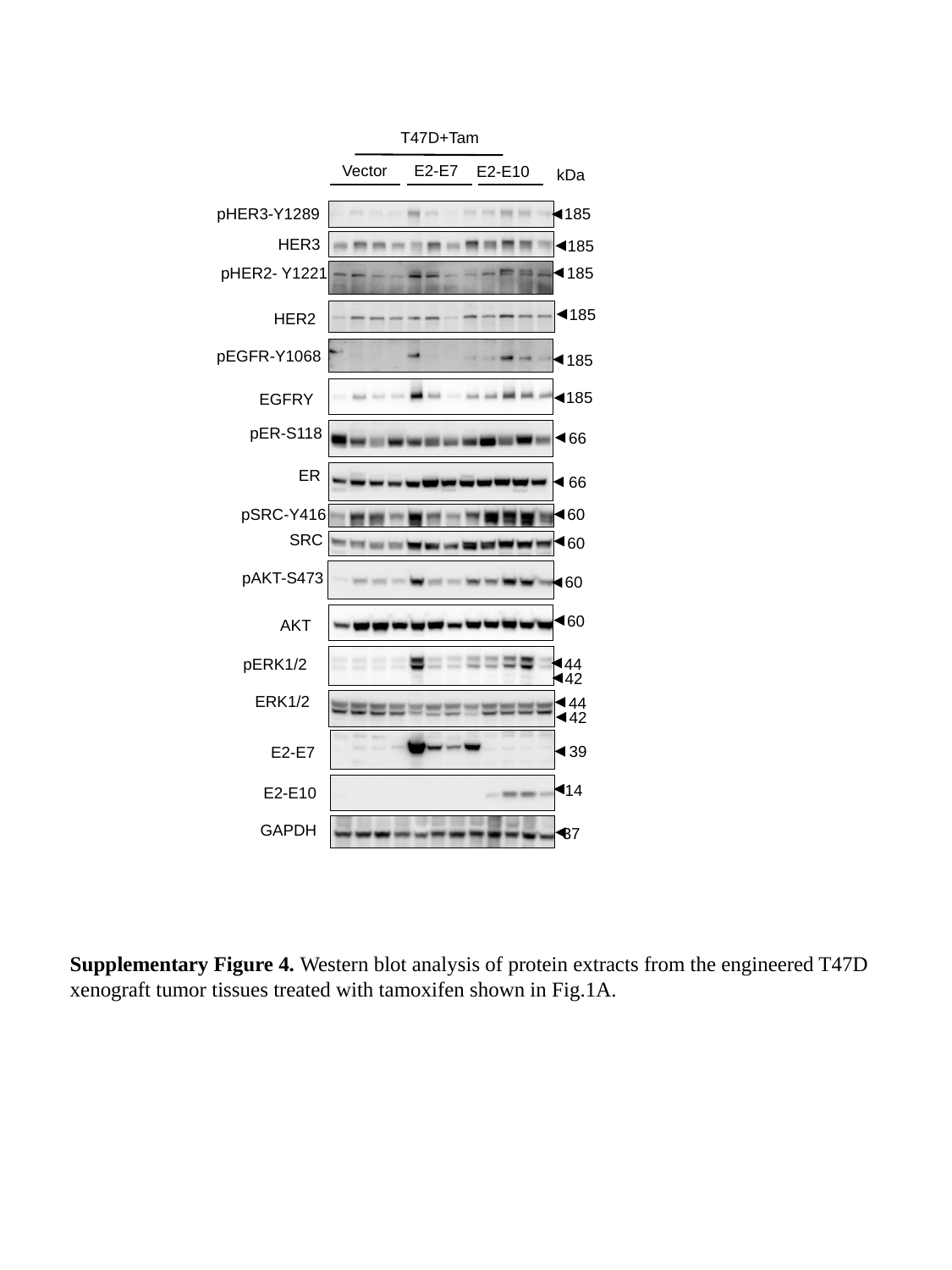

T47D+Tam
Vector
E2-E7
E2-E10
kDa
pHER3-Y1289
185
HER3
185
pHER2- Y1221
185
185
HER2
pEGFR-Y1068
185
185
EGFRY
pER-S118
 66
 66
pSRC-Y416
 60
SRC
 60
pAKT-S473
 60
 60
AKT
 44
pERK1/2
 42
ERK1/2
 44
 42
 39
E2-E7
 14
E2-E10
GAPDH
 37
ER
Supplementary Figure 4. Western blot analysis of protein extracts from the engineered T47D xenograft tumor tissues treated with tamoxifen shown in Fig.1A.
