## Additional File 5 for "Therapeutic role of recurrent ESR1-CCDC170 gene fusions in breast cancer endocrine resistance"

### Slide 1
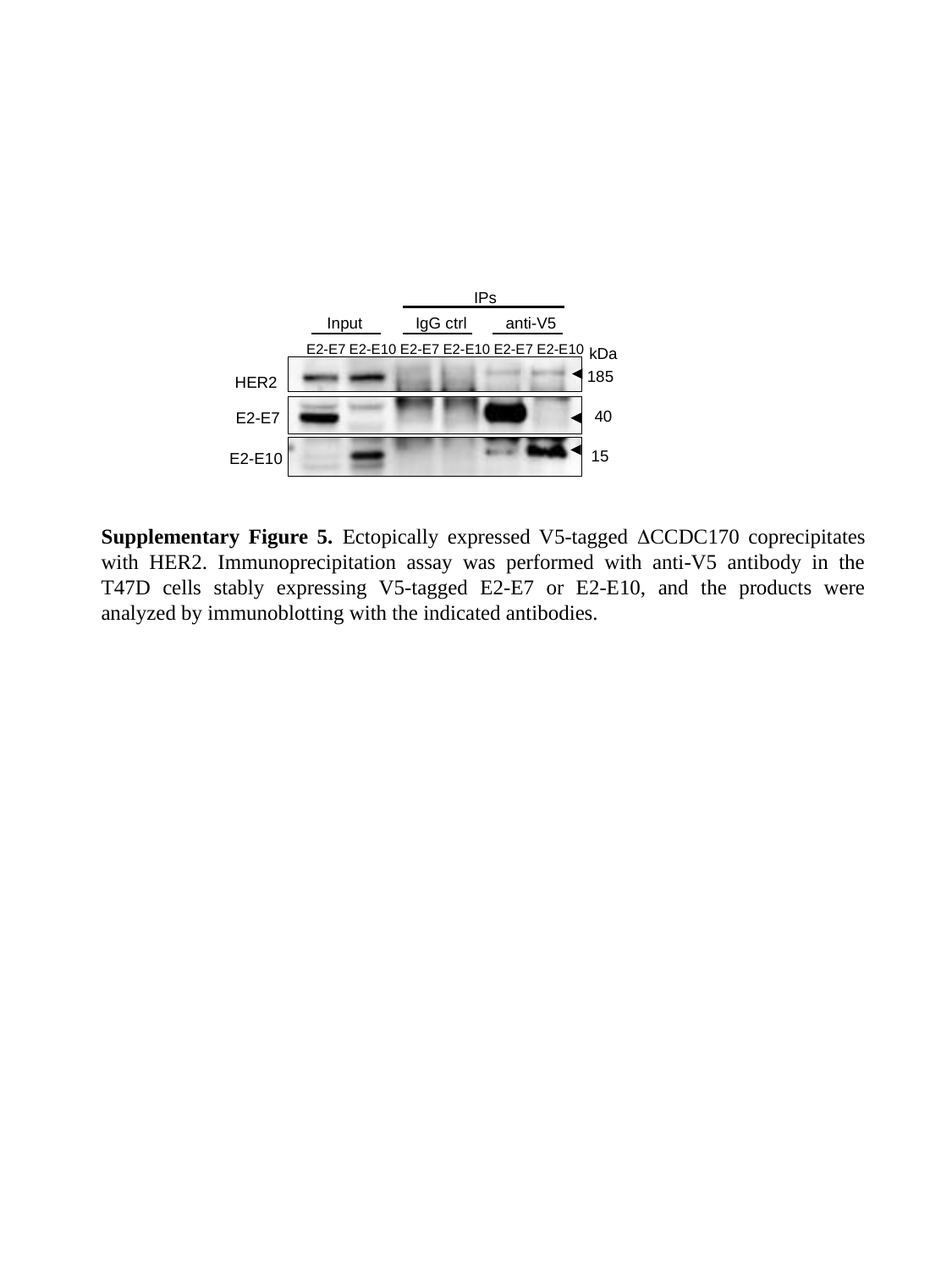

IPs
Input
IgG ctrl
anti-V5
E2-E7
E2-E10
E2-E7
E2-E10
E2-E7
E2-E10
HER2
E2-E7
E2-E10
185
40
15
kDa
Supplementary Figure 5. Ectopically expressed V5-tagged ΔCCDC170 coprecipitates with HER2. Immunoprecipitation assay was performed with anti-V5 antibody in the T47D cells stably expressing V5-tagged E2-E7 or E2-E10, and the products were analyzed by immunoblotting with the indicated antibodies.
