## Additional File 6 for "Therapeutic role of recurrent ESR1-CCDC170 gene fusions in breast cancer endocrine resistance"

### Slide 1
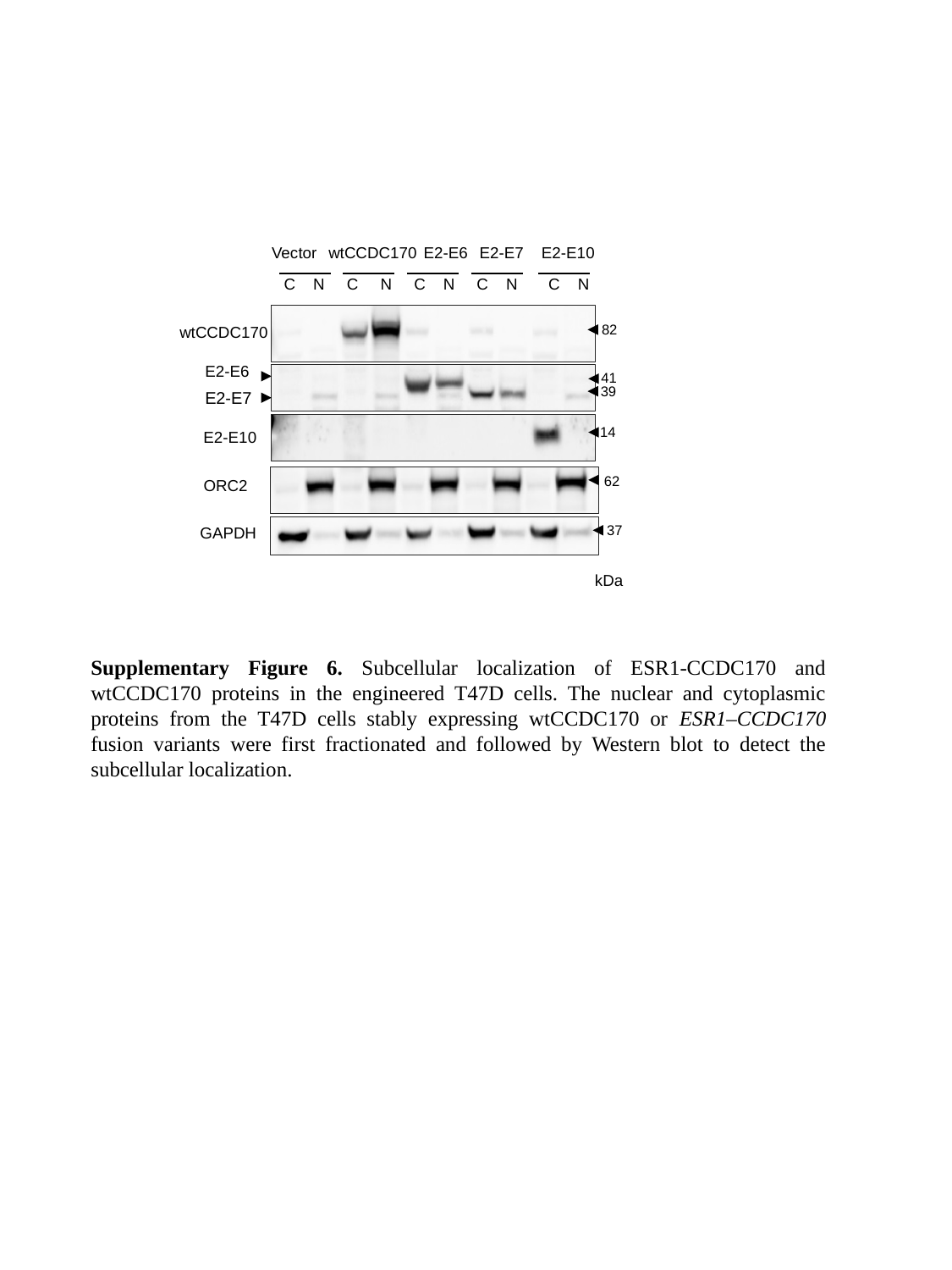

Vector
wtCCDC170
E2-E6
E2-E7
E2-E10
C N C N C N C N C N
82
wtCCDC170
E2-E6
41
39
E2-E7
14
E2-E10
62
ORC2
37
GAPDH
kDa
Supplementary Figure 6. Subcellular localization of ESR1-CCDC170 and wtCCDC170 proteins in the engineered T47D cells. The nuclear and cytoplasmic proteins from the T47D cells stably expressing wtCCDC170 or ESR1–CCDC170 fusion variants were first fractionated and followed by Western blot to detect the subcellular localization.
