## Additional File 7 for "Therapeutic role of recurrent ESR1-CCDC170 gene fusions in breast cancer endocrine resistance"

### Slide 1
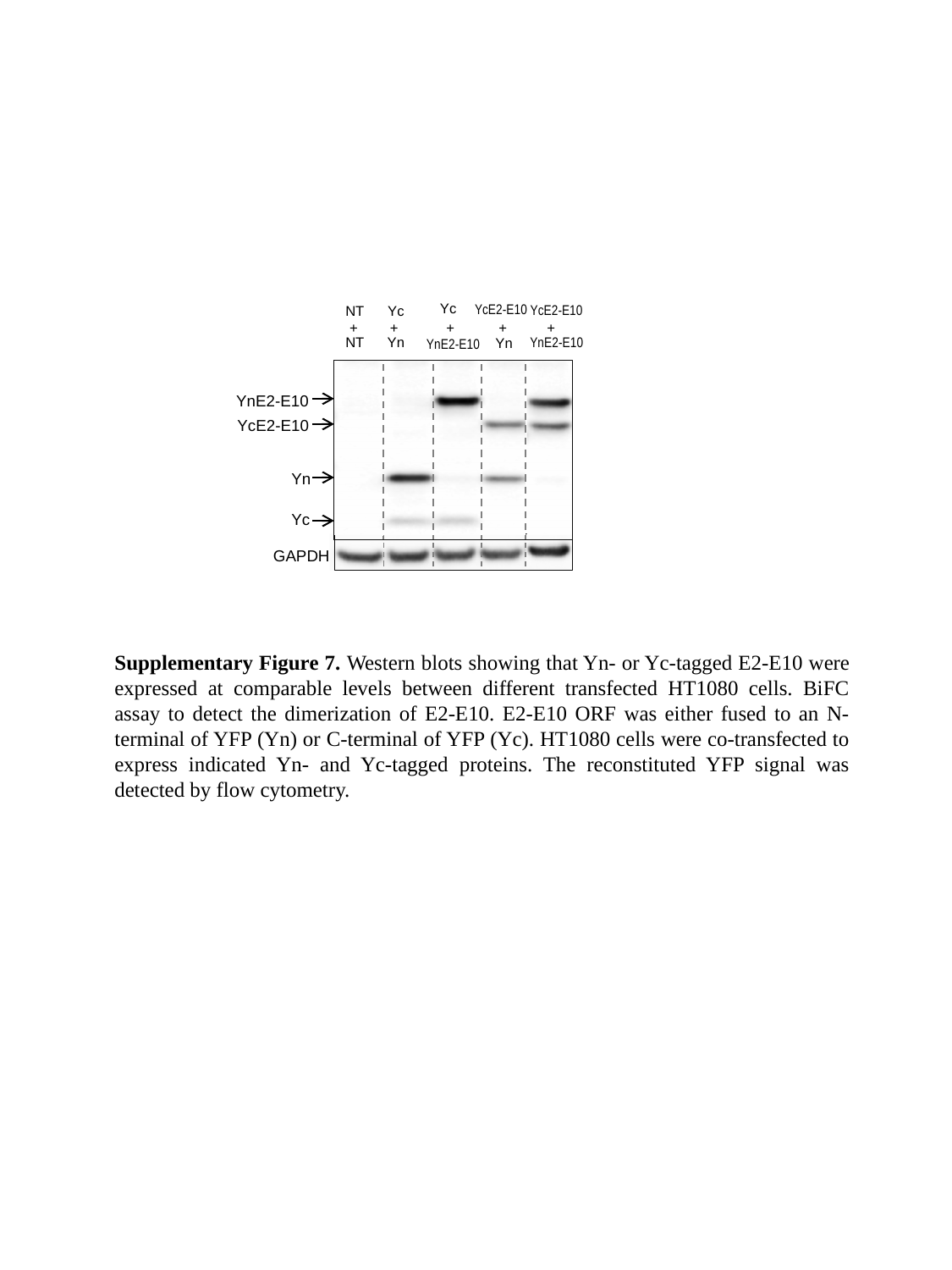

Yc
YcE2-E10
NT
Yc
+ + + + +
NT
Yn
YnE2-E10
YnE2-E10
YcE2-E10
Yn
Yc
GAPDH
YnE2-E10
Yn
YcE2-E10
Supplementary Figure 7. Western blots showing that Yn- or Yc-tagged E2-E10 were expressed at comparable levels between different transfected HT1080 cells. BiFC assay to detect the dimerization of E2-E10. E2-E10 ORF was either fused to an N-terminal of YFP (Yn) or C-terminal of YFP (Yc). HT1080 cells were co-transfected to express indicated Yn- and Yc-tagged proteins. The reconstituted YFP signal was detected by flow cytometry.
