## Additional File 8 for "Therapeutic role of recurrent ESR1-CCDC170 gene fusions in breast cancer endocrine resistance"

### Slide 1
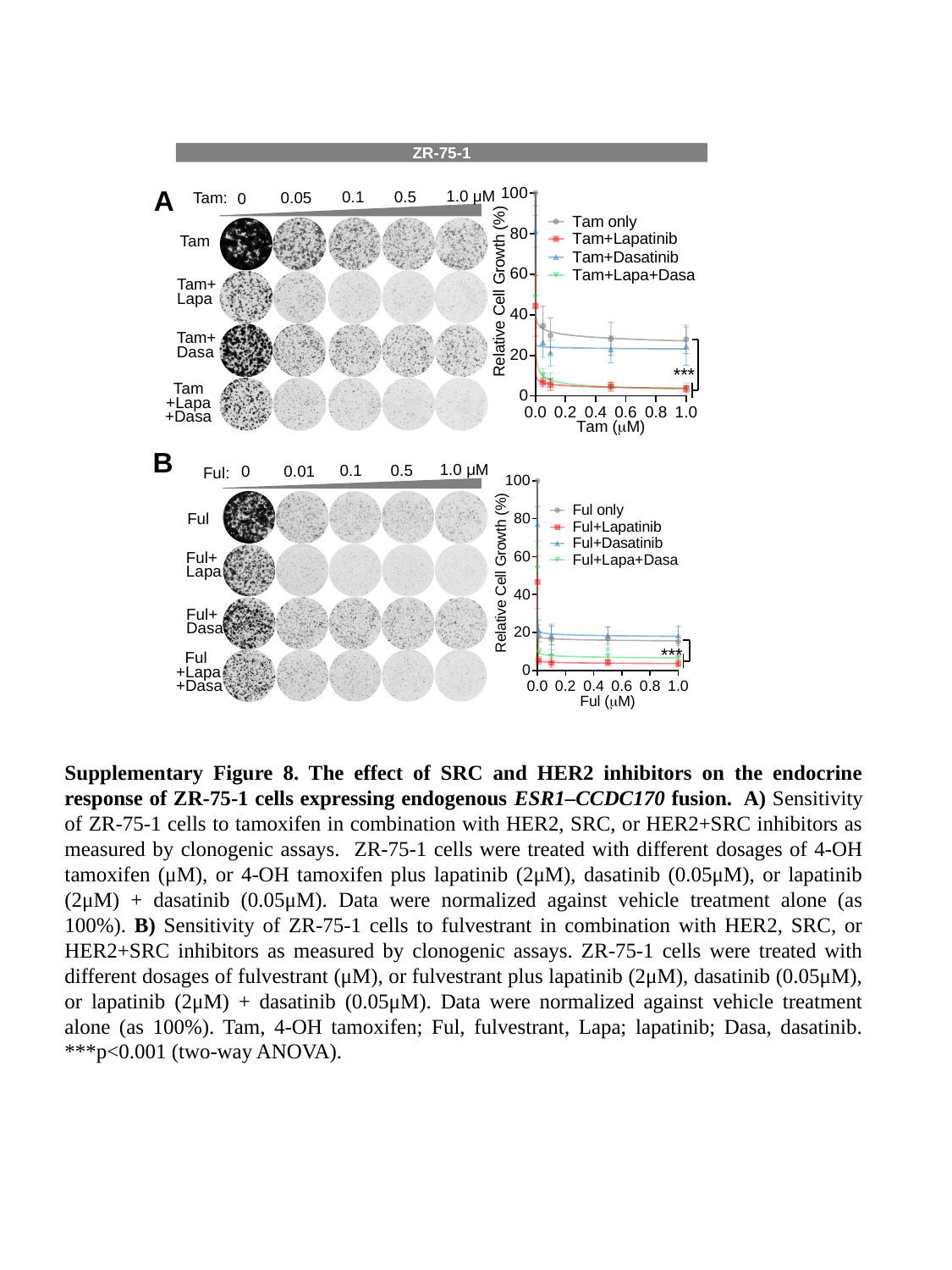

ZR-75-1
A
1.0 μM
0.5
0.1
0.05
Tam:
0
Tam
Tam+
Lapa
Tam+
Dasa
Tam
+Lapa
+Dasa
***
B
1.0 μM
0.1
0.5
0
0.01
Ful:
Ful
Ful+
Lapa
Ful+
Dasa
 Ful
+Lapa
+Dasa
***
Supplementary Figure 8. The effect of SRC and HER2 inhibitors on the endocrine response of ZR-75-1 cells expressing endogenous ESR1–CCDC170 fusion. A) Sensitivity of ZR-75-1 cells to tamoxifen in combination with HER2, SRC, or HER2+SRC inhibitors as measured by clonogenic assays. ZR-75-1 cells were treated with different dosages of 4-OH tamoxifen (μM), or 4-OH tamoxifen plus lapatinib (2μM), dasatinib (0.05μM), or lapatinib (2μM) + dasatinib (0.05μM). Data were normalized against vehicle treatment alone (as 100%). B) Sensitivity of ZR-75-1 cells to fulvestrant in combination with HER2, SRC, or HER2+SRC inhibitors as measured by clonogenic assays. ZR-75-1 cells were treated with different dosages of fulvestrant (μM), or fulvestrant plus lapatinib (2μM), dasatinib (0.05μM), or lapatinib (2μM) + dasatinib (0.05μM). Data were normalized against vehicle treatment alone (as 100%). Tam, 4-OH tamoxifen; Ful, fulvestrant, Lapa; lapatinib; Dasa, dasatinib. ***p<0.001 (two-way ANOVA).
