## Additional File 9 for "Therapeutic role of recurrent ESR1-CCDC170 gene fusions in breast cancer endocrine resistance"

### Slide 1
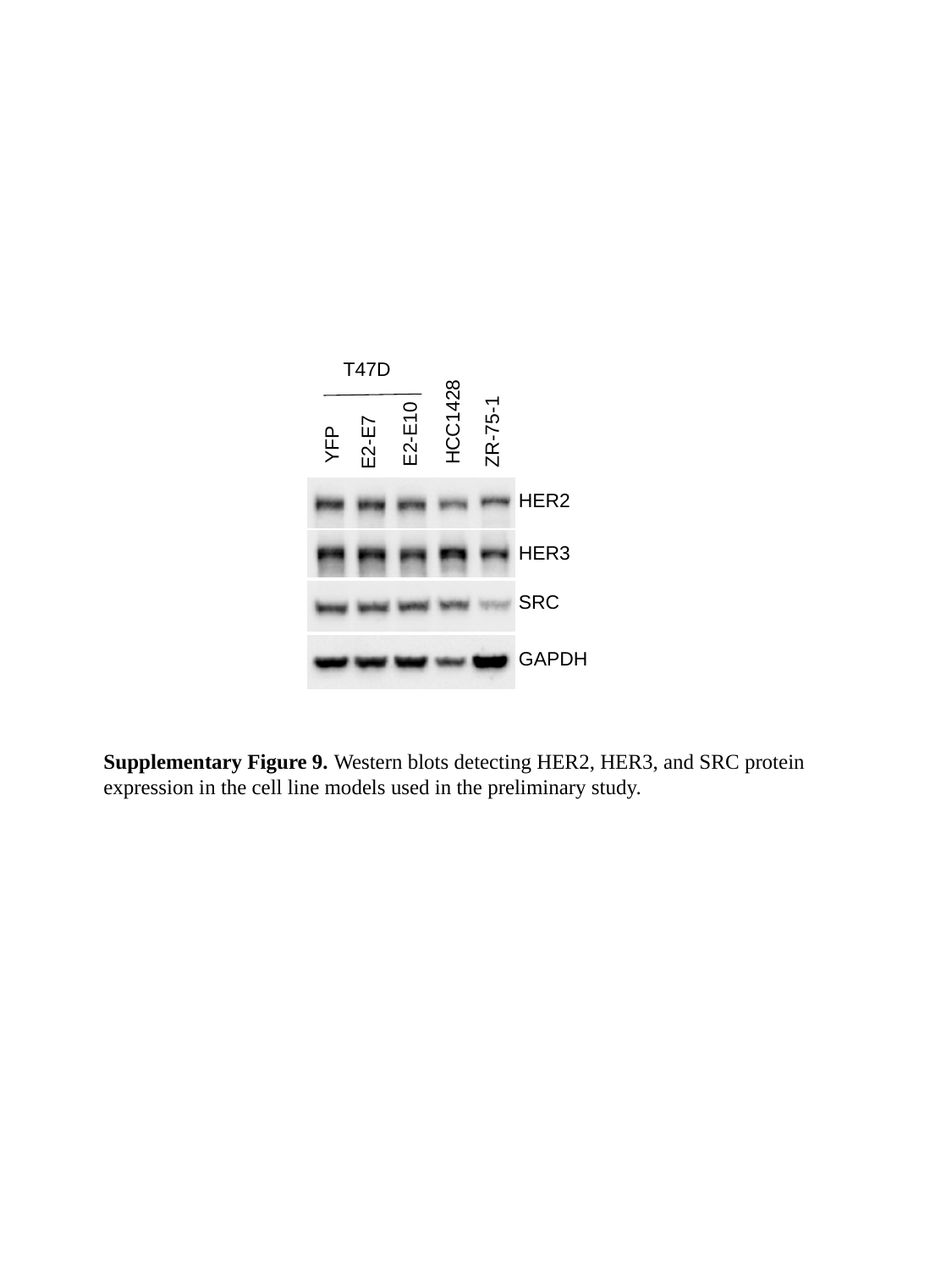

T47D
HCC1428
ZR-75-1
E2-E10
E2-E7
YFP
HER2
HER3
SRC
GAPDH
Supplementary Figure 9. Western blots detecting HER2, HER3, and SRC protein expression in the cell line models used in the preliminary study.
